# Nurr1 orchestrates claustrum development and functionality

**DOI:** 10.1101/2024.12.31.630879

**Authors:** Kuo Yan, Andrew Newman, Pauline Lange, Susanne Mueller, Marco Foddis, Maron Mantwill, Carsten Finke, Stefan Paul Koch, Philipp Boehm-Sturm, Penghui Deng, Melissa Long, Victor Tarabykin

## Abstract

Claustrum is the part of the forebrain known to be most widely interconnected tissue with almost all global regions. It is believed to be involved in coordinating multiple cognitive behaviors including consciousness formation. However, little is known about the molecular mechanisms underlying its development and involvement in behavioral control. Here we show that Nurr1 (Nr4a2) is the key transcription factor orchestrating claustral morphogenesis, functional connectivity and thus behaviors. Nurr1 deficient claustral cells aberrantly migrate into insular cortex, shaping claustrum into an abnormal wing-like structure, and ectopically turn on insular corticex genetic program. Accordingly, functional connectivity of the claustrum is not properly formed and relevant behaviors are dysregulated in Nurr1 deficient mice. We show that Nurr1 regulates claustral neuron positioning by suppressing G-protein signaling.

## Introduction

The claustrum (CLA), a sheet-like gray matter structure resident between striatum and insular cortex in mice, exists in most vertebrate species ranging from birds, reptiles, rodents to human^1–4^. Little attention has been paid to this thin structure since being identified 200 years ago until Crick and Koch proposed that CLA be the processing hub of sensory information from multiple cerebral sources to integrate coherent consciousness^5^. Though whether CLA is the core hardware of consciousness is still uncertain to date, it is indeed intensively interconnected to almost all global subareas^4, 6–8^, empowering its involvement in diverse cognitive activities, including but not limited to stress responses, attention, salient sensation, sensorimotor cross-modal selection, sleep and pain perception^9–14^. The insights into nerve circuitry underlying these physiological functions are accumulating due to recent advancement of optogenetics^9, 15,16^. However, very little has been elucidated about the molecular mechanisms governing CLA development and CLA-dependent cognitive behaviors yet.

Nurr1 (Nr4a2) is a transcription factor that contains a single zinc-finger DNA-binding domain. It is not only renowned to promote dopaminergic neurogenesis as well as dopamine production in substantia nigra^17, 18^, but also the most frequently used marker for CLA glutamatergic neurons^1, 19, 20^. Nurr1 expression in CLA neurons is already detectable as early as E13.5 and lasts throughout life in mice (*Allen brain atlas*), but its role in CLA remains largely unknown. Here we demonstrate novel evidence that Nurr1’s transcription activity is indispensable to orchestrate claustral morphogenesis, cell identity determination, functional connectivity and CLA-dependent cognitive behaviors.

## Materials and Methods

The details of “*Materials and Methods*” are included in supplement.

## Results

### Transcription activity of Nurr1 is necessary for claustrum morphogenesis

In order to characterize Nurr1’s roles in forebrain development, we generated Nurr1 conditionally deficient mice regulated by EmxCre (Nurr1^flox/flox^;EmxCre^Cre/Wt^, Nurr1 deficient mice throughout text unless otherwise noted), where the floxed exon 3 of Nurr1’s transcript containing the start codon and DNA-binding domain was selectively depleted in dorsal telencephalon at ∼E10 by Cre recombinase under the control of Emx1 promoter^21^. The genetic design of Nurr1^flox/flox^ mouse model entitles a likelihood to trace Nurr1-expressing cells absent of its transcription regulatory activity in Cre-present brains in that a C-terminal (Cterm) truncated open reading frame of Nurr1 consistent with the original transcript under control of the same Nurr1 promoter might be still present while recombined^18^. To testify this, we procured antibodies to target N-terminal (Nterm) or Cterm portions of Nurr1 protein, as well as *in situ* hybridization (ISH) probes to target Nurr1 5’- or 3’-terminal transcripts (5’end or 3’end), respectively (Fig. 1A). While ISH by the 5’end probe verified Nurr1 expression depletion in mutant brains (Fig. S1A), Nurr1 3’end mRNA was still detectable in CLA, subplate (Sp), dorsal endopiriform nuclei (dEn), subiculum (Sb) and cortical plate (CP) (Fig. S1B), similar to Nurr1 expression in controls. Likewise, immunofluorescence (IF) using Nurr1 Nterm and Cterm antibodies confirmed the same results (Fig. S1C). Notably, the truncated Nurr1-Cterm polypeptide lost nuclear localization (Fig. S1C) due to depletion of nuclear localization signal peptide^22^, indicating its inability to regulate transcription.

**Figure 1.**
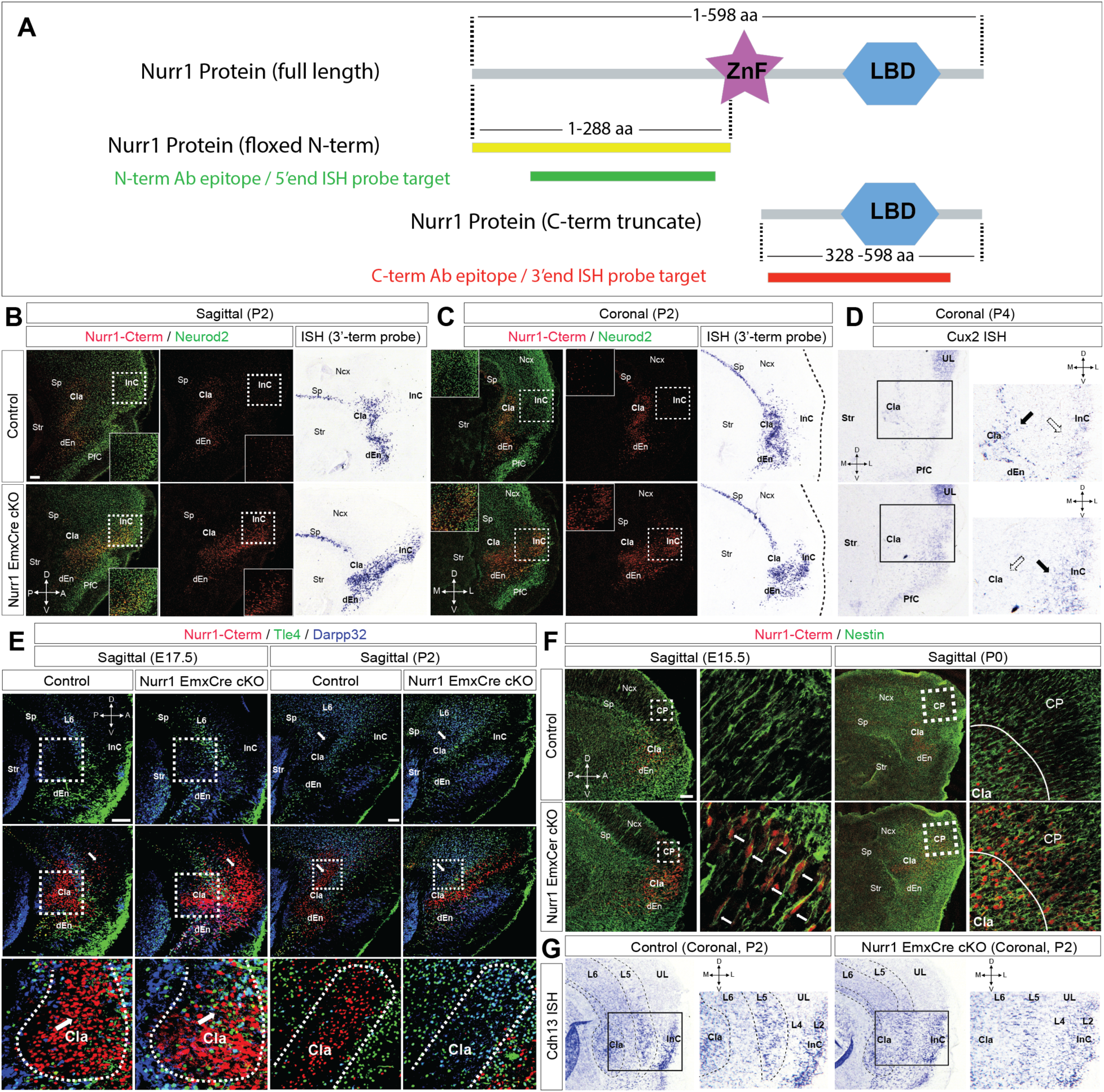
Nurr1 is a key regulator for claustrum morphogenesis. **(A)** Illustration of Nurr1 molecular structure. The full-length (FL) Nurr1 protein consists of a DNA-binding zinc finger (ZnF) domain and a ligand-binding domain (LBD). The coding sequence of N-terminal region (Nterm, yellow, 1 – 288 amino acids (aa)) of Nurr1 protein is floxed (Kadkhodaei, et al., 2009). The truncated C-terminal portion (Cterm, red, 328 – 598 aa) of Nurr1 protein in the same original open reading frame loses ZnF domain after Cre-mediated recombination. The protein epitopes targeted by the antibodies (ab) or the transcript sequences targeted by the ISH probes are represented in green (Nterm ab or 5’end probe) and red (Cterm ab or 3’end probe), respectively. **(B, C)** Immunofluorescence (IF) staining for Nurr1-Cterm (red) and Neurod2 (green) on sagittal **(B)** and coronal (**C**) sections of control and EmxCre+ Nurr1 deficient (cKO) brains at P2. Nurr1/Neurod2 double positive (Nurr1+/Neurod2+) claustral (CLA) neurons are mostly positioned between striatum (str) and insular cortex (InC), however, a huge population of Nurr1 deficient cells over-migrate in InC territory. ISH using the Nurr1-3’end probe on sagittal and coronal sections show the same migration deficits of Nurr1 deficient CLA cells as the IF data. The framed images show magnification views of the dotted-line boxed areas. Sp, subplate; dEn, dorsal endopiriform nucleus; PfC, piriform cortex, Ncx, neocortex. D, dorsal; A, anterior; V, ventral; P, posterior, M, medial; L, lateral. Scale bar: 200 µm (the same in this figure). **(D)** ISH using the Cux2 probe on coronal sections of control and Nurr1 cKO brains at P2. Cux2 is strongly expressed in a subpopulation of CLA neurons (solid arrowhead) and weakly expressed in InC (hollow arrowhead), but its expression was ectopically detected in the InC of Nurr1 cKO brains but no longer in CLA, implying that Cux2+ CLA neurons over-migrate into InC. The enlarged images show magnification views of the boxed areas (the same below in all figures). UL, upper layers of Ncx. **(E)** IF for Nurr1-Cterm (red), Tle4 (green) and Darpp32 (blue) on sagittal sections of control and EmxCre+ mutant brains at E17.5 and P2. CLA neurons are already largely ceased in the area surrounded by Tle4+/Darpp32+ deeper layer (DL) neurons at E17.5, and these DL neurons delineate the border for Nurr1+ CLA neurons at P2 in control brains. However, the majority of Nurr1 deficient CLA cells already migrate across Tle4+/Darpp32+ DL neurons at E17.5 and are eventually located in InC anterior to DL neurons at P2. The original CLA area is filled with Tle4+/Darpp32+ DL neurons in Nurr1 deficient brains. The arrowheads indicate CLA areas in control and cKO brains. The dotted lines delineate CLA territories from DL neurons. **(F)** IF for Nurr1-Cterm (red) and Nestin (green) on sagittal sections of control and EmxCre+ mutant brains at E15.5 and P0. Nurr1 deficient CLA neurons always keep radial migration along Nestin+ radial glia. The lines in P0 images delineate the borders between CLA and Ncx. CP, cortical plate. **(G)** ISH using the Cdh13 probe on coronal sections of control and EmxCre+ mutant brains at P2. The laminated expression of Cdh13 depicts the cortical cytoarchitecture in control brains by displaying its stronger expression in CLA, layer 5 (L5) and L2 of Ncx, but weaker expression in L6 and L4 along medio-lateral axis. However, the clear cytoarchitecture is disrupted by migrating Nurr1 deficient cells.

The general morphology of Nurr1 deficient cortex are basically normal (Fig. S2A-B), consistent with previous reports^17, 18^. We then focused on the Nurr1-expressing structures and found that Sp appeared normal in morphology in Nurr1 deficient brains using ISH for Sp markers - CTGF and S100a10^23, 24^ (Fig. S2C-D). ISH for Zbtb20, a hippocampal marker^25, 26^, in developing and postnatal brains suggested that the formation of perisubicular complexes was also normal in Nurr1 deficient brains (Fig. S2E-F). Additionally, co-IF for Nurr1 and Zbtb20 depicted that Nurr1-positive (Nurr1+) Sb delineated borders to separate neocortical cingulate cortex from hippocampal distal CA1, which maintained comparable in control and mutant brains (Fig. S2G). In contrast, both IF and ISH data demonstrated that Nurr1 deficient CLA cells detached from Sp and invade insular cortex (InC) in both sagittal and coronal views (Fig. 1B-C). To further confirm the mislocalization of Nurr1 deficient CLA cells, we performed ISH for Cux2, another CLA cell marker^20^, and found Cux2+ CLA cells become ectopically present in InC of Nurr1 deficient brains (Fig. 1D). Consequently, the CLA was reshaped from a crescent-like into a wing-like structure (Fig. S3A). We next asked if the ectopic CLA cells in InC of Nurr1 deficient brains were due to over-production. We quantified Nurr1-Cterm+ cells in claustro-insular region and found its number in Nurr1 deficient brains was similar with that in controls (Fig. S3B). Furthermore, few cleaved-Caspase3+ cells were detected in the claustro-insular regions of control and mutant brains (Fig. S3C-D). Nurr1 thus does not play a role in CLA neuron productivity nor survival.

We then questioned what happened to the original CLA territory when Nurr1 deficient cells moved away. IF for Nurr1-Cterm and Tle4/Darpp32 delineated CLA cells from neighboring InC deeper layer (DL) neurons in control brains, depicting normal CLA cells were bilaterally restricted in an area between stratum and InC DL neurons. However, Nurr1 deficient CLA cells had largely bypassed InC DL cells since E17.5 (Fig. 1E). Intriguingly, Tle4/Darpp32-expressing DL cells, which were not found in the control CLA area, became abundant in the original CLA territory of postnatal mutant brains (Fig. 1E), an observation further strengthened by ISH for Fezf2 and Nfib (Fig. S4). It has been proposed Nurr1+ neurons can migrate radially and tangentially to CLA and neocortex^19^, we next wanted to study the migration mode of Nurr1 deficient CLA cells. IF for Nurr1-Cterm and Nestin showed Nurr1- Cterm+ CLA cells in mutant brains constitutively migrated towards pia along radial glia scaffold throughout cortical neurogenesis (Fig. 1F), thus spoiling the laminated organization of claustro-insular tissues (Fig. 1G). Collectively, Nurr1 orchestrates multiple aspects of CLA morphogenesis, such as positioning and interaction with adjacent tissues, to make CLA such distinct cytoarchitecture.

Another key question is whether Nurr1 determines CLA cell positioning pre- or postmitotically. To answer this, we generated Nurr1^flox/flox^;NexCre^Cre/Wt^ mice, in which Nurr1 was selectively inactivated in cortical postmitotic compartment^27^. It turned out that Nurr1 NexCre mutant brains fully recapitulated the morphological phenotypes as in EmxCre mutant brains (Fig. S5A-B), suggesting Nurr1’s role in postmitotic regulation.

### Nurr1 controls claustral neuron specification postmitotically

We next desired to find out if Nurr1 also played a role in CLA cell specification. To this end, we first performed ISH for several CLA marker genes, such as NtnG2, Gnb4 and Gng2^20, 28^, and found all these genes were significantly downregulated in CLA (and/or dEn neurons) of both EmxCre (Fig. 2A-C and 2A’-C’) and NexCre (Fig. S5C-E) mutant brains. Furthermore, single cell mRNA sequencing (scmRNA-seq) of claustro-insular tissues revealed a large spectrum of misregulated genes in Nurr1 deficient brains (see below, Fig. 5, S8 and S9), among which, Rgs20, for instance, was selectively attenuated in CLA, but not in Sp (Fig. 2D-D’). These results strongly indicate the specific genetic profile of CLA is altered due to Nurr1 deficiency. Secondly, we tested if Nurr1 deficient CLA cells activated InC gene expression thereupon their ectopic localization. Cyp26b1 is a widely accepted InC marker^29,30^. We quantified the proportion of Nurr1/Cyp26b1 double positive (Nurr1+/Cyp26b1+) cells relative to total Nurr1+ cells in claustro-insular region. While few Nurr1+/Cyp26b1+ cells were detected in controls, a considerably increased portion of Nurr1-Cterm+ cells (by ∼9.70 ± 1.64 folds, *p* = 0.00010, ***) in the InC of EmxCre mutant brains switched on Cyp26b1 expression (Fig. 2E). By the same token, we quantified the proportions of Nurr1+/Neurod1+ and Nurr1+/Rorβ+ cells in claustro-insular region as Neurod1 and Rorβ are also preferentially expressed in InC cells but not in CLA *(Allen brain atlas*). The proportions of Nurr1+/Neurod1+ and Nurr1+/Rorβ+ cells were increased by ∼4.49 ± 0.36 folds (*p* < 0.0001, ****) and by ∼19.92 ± 0.72 folds (*p* < 0.0001, ****) in EmxCre mutant brains relative to those of control brains, respectively (Fig. 2F, 2G). Furthermore, the expression levels of these InC enriched genes in Nurr1 NexCre mutant brains were elevated to the comparable levels as those in EmxCre mutant brains (Fig. 2E-G). These results suggest Nurr1 deficient CLA cells tend to convert their genetic program to InC cells’ postmitotically.

**Figure 2.**
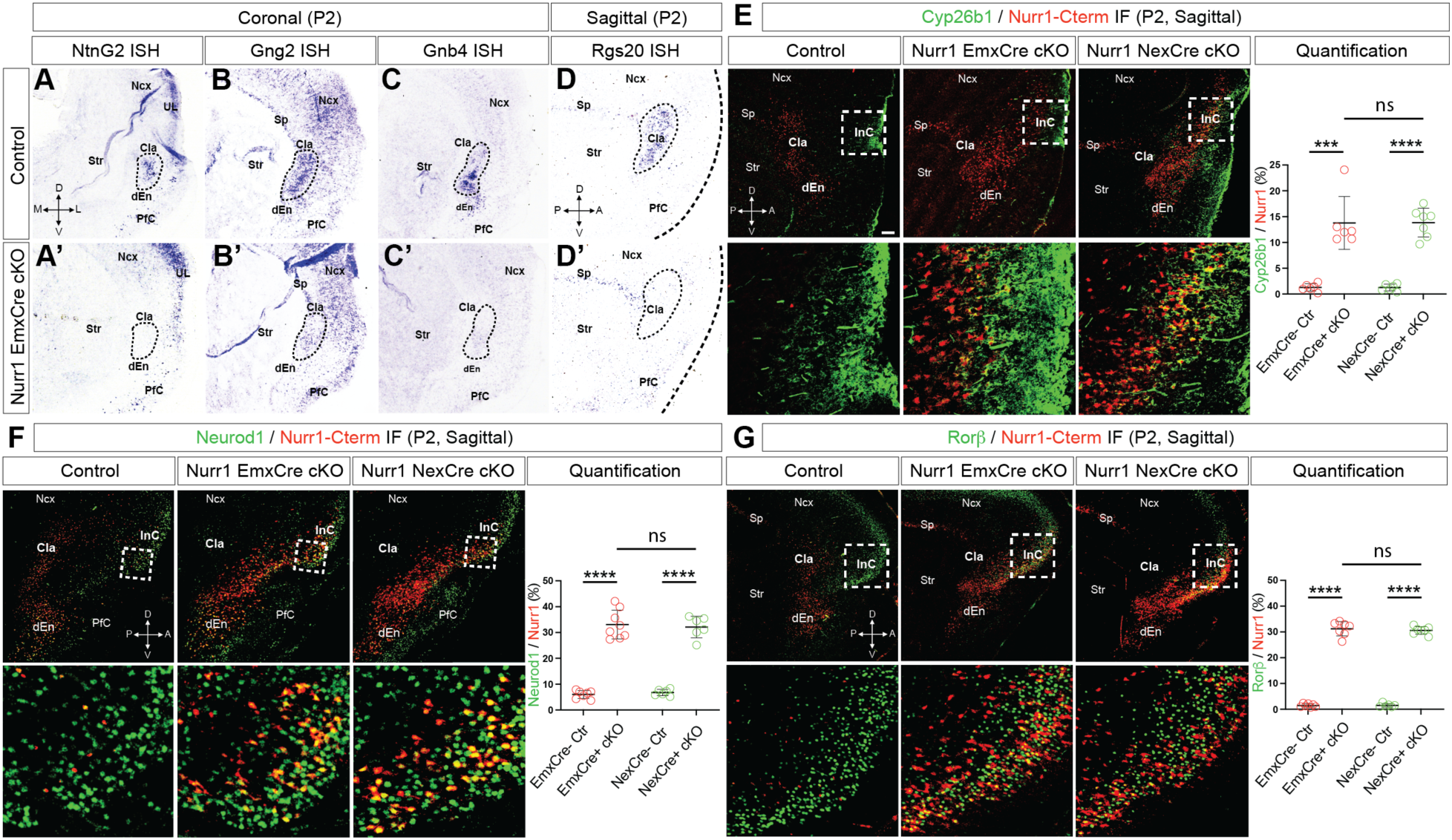
Nurr1 is essential to determine claustral cell fate. (A-A’ *–* D-D’) ISH using the NtnG2, Gng2, Gnb4 and Rgs20 probes on brain sections of control and EmxCre+ mutant brains at P2. NtnG2 is expressed in CLA, UL and PfC in control brains but selectively downregulated in CLA of Nurr1 deficient brains (**A-A’**). Gng2 expression is widely detectable in CLA, Sp, PfC and Ncx in control brains (**B**), whereas its expression is severely reduced in ClA but not in surrounding areas of mutant brains (**B’**). Gnb4 is specifically expressed in CLA and dEn neurons in control brains (**C**) but becomes rarely detectable in Nurr1 deficient brains (**C’**). (**D-D’**) ISH using the Rgs20 probe on sagittal sections of control and EmxCre+ mutant brains at P2. Rgs20 is normally expressed in Sp and CLA but selectively absent in CLA in mutant brains. The dotted-line circles in (**A-A’ *–* D-D’**) indicate CLA areas. **(E *–* G)** IF for Nurr1-Cterm with Cyp26b1 (**E**), Neurod1 (**F**) and Rorβ (**G**) on sagittal sections of control, EmxCre+ and NexCre+ Nurr1 deficient brains at P2. Nurr1+/Cyp26b1+ cells are rarely detectable in control brains but significantly increased in both EmxCre+ and NexCre+ mutant brains (**E**). The proportion of Nurr1+/Cyp26b1+ cells in EmxCre+ mutant brains (≈13.77%, n = 6) is increased by ∼9.70 ± 1.64 folds (*p* = 0.00010, ***) relative to controls (≈1.29%, n = 6). The proportion in NexCre+ cKO brains (≈13.85%, n = 7) is increased by ∼9.95 ± 0.92 folds (*p* < 0.0001, ****) relative to controls (≈1.27%, n = 6), to the similar level with EmxCre+ mutant brains (*p* = 0.97, ns). The proportions of Nurr1+/Neurod1+ cells in EmxCre+ and NexCre+ mutant brains are also elevated to similar levels (*p* = 0.72, ns): the proportion in EmxCre+ mutant brains (≈33.11%, n = 8) is increased by ∼4.49 ± 0.36 folds (p < 0.0001, ****) relative to controls (≈ 6.03%, n = 7), and the proportion in NexCre+ cKO brains (≈32.09%, n = 6) is increased by ∼3.73 ± 0.26 folds (*p* < 0.0001, ****) relative to controls (≈6.78%, n = 6). Likewise, the proportions of Nurr1+/Rorβ+ cells in EmxCre+ and NexCre+ mutant brains are elevated to similar levels (*p* = 0.60, ns): the proportion in EmxCre+ mutant brains (≈31.23%, n = 7) is increased by ∼19.92 ± 0.72 folds (*p* < 0.0001, ****) relative to controls (≈1.49%, n = 7), and the proportion in NexCre+ cKO brains (≈30.59%, n = 7) is increased by ∼19.02 ± 0.44 folds (*p* < 0.0001, ****) relative to controls (≈1.53%, n = 6). The statistics data was analyzed by two-tailed *Student’s t*-test. Scale bar: 200 µm.

Axonal projection pattern is one of the major aspects of neuron subtype identity^31^. To test if the axonal connections of Nurr1 deficient neurons are also altered accordingly, we performed retrograde axonal tracing by placing DiI crystals into two target regions – primary motor cortex (PMC, telencephalic) and hypothalamus (HT, diencephalic), which were reported to be connected intensively with InC, but very weakly with CLA^6, 32–36^. While efferent neurons to PMC were rarely rooted in CLA in control brains, Nurr1 deficient cell somas in InC frequently colocalized with DiI retrograde signals, supported by that the ratio of Nurr1+/DiI+ cells in Nurr1 deficient brains was increased by ∼28.58 ± 4.53 folds compared with that in controls (Fig. S6A). Likewise, the ratio of Nurr1-Cterm+/DiI+ cells in mutant brains was approximately triple of that in controls in the scenario of HT retrograde labeling (Fig. S6B). Taken together, Nurr1 plays a decisive role in CLA cell identity establishment.

### Functional connectivity associated with claustrum is disrupted in Nurr1 deficient mice

Given Nurr1’s role in axonal wiring and global interconnection of CLA neurons, we thus studied if connectivity mapping in Nurr1 deficient mice was altered using resting state functional magnetic resonance imaging (rs-fMRI). We first analyzed the functional connectivity (FC) of 17 cerebral networks (see “*Materials and Methods*”) that had been reported to be connected to CLA^6, 35, 36^, and found the amygdala and anterior cingulate cortex (ACC) networks were the most severely affected in Nurr1 deficient brains (Fig. 3A), supported by their clusters of significantly lower network activity (i.e. independent component weight) than those of control brains (Fig. 3B, 3C). In order to find out if the compromised plasticity in these networks is specifically correlated to CLA, we employed the method of seed-based differential mapping according to the *Allen brain atlas*^8^. When both hemispheric CLA were seeded as the regions of interest (ROI), synchronous activity was strongly detected in amygdala and ACC networks in control brains, but considerably weakened in Nurr1 deficient brains (Fig. 3D-D’), suggesting the impaired FC in these networks of Nurr1 deficient brains is linked with CLA abnormality. Analysis based on single hemispheric CLA seed did not only depict similar results (sagittal plane, Fig. 3E-E’), but also revealed that CLA was correlated to interhemispheric FC of amygdala and ACC networks, however, both were reduced in Nurr1 deficient brains (coronal plane, Fig. 3E-E’).

**Figure 3.**
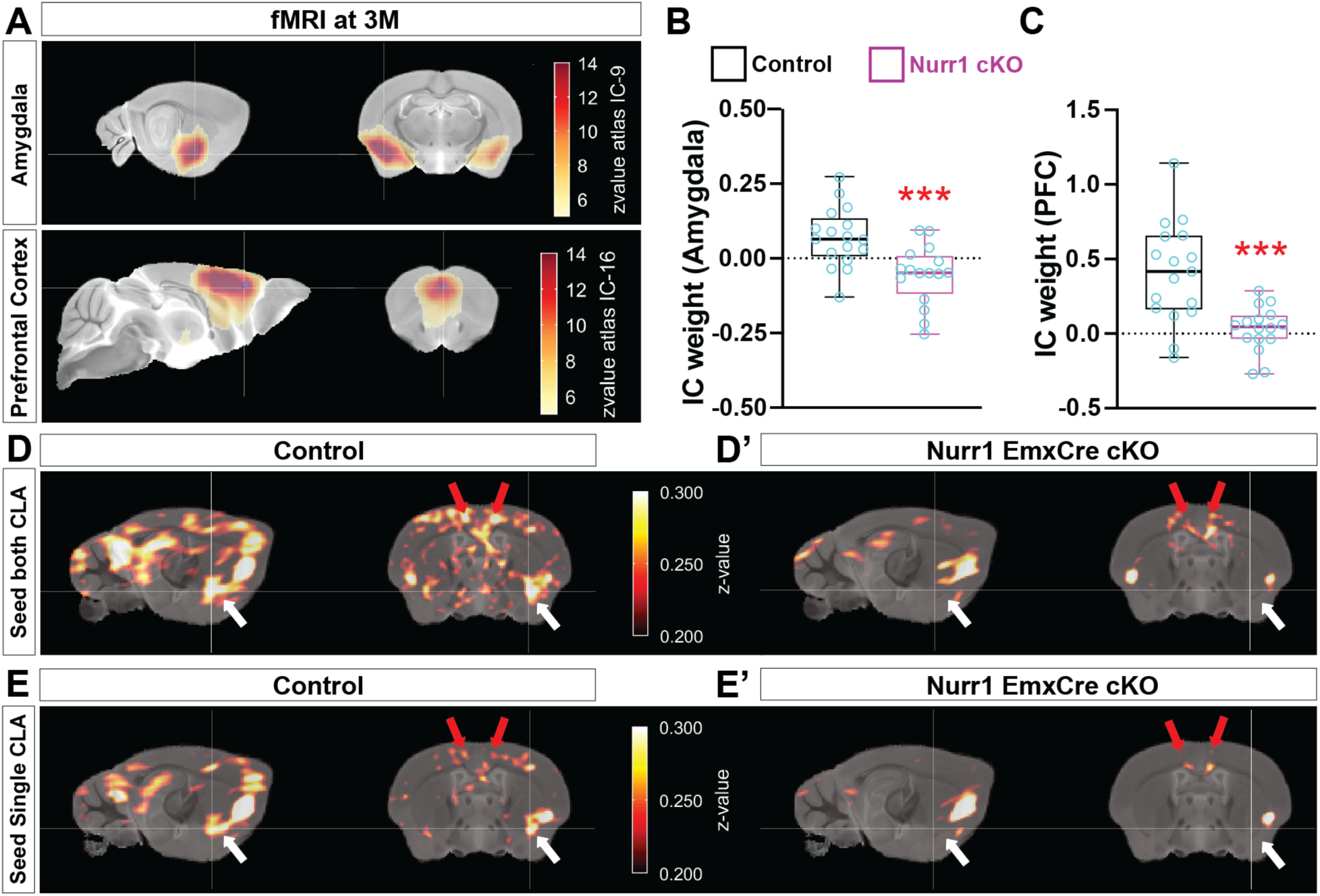
Connectivity networks associated with claustrum are impaired in Nurr1 deficient brains. (A –. **C)** Resting state fMRI (rs-fMRI) was used to examine functional connectivity of 17 networks of control and Nurr1 deficient brains. (**A**) Blood-oxygen-level dependent (BOLD) signal intensities in amygdala and anterior cingulate cortex (ACC) networks (red z-score overlaid on top of grayscale anatomical MRI template) of control brains are positively over threshold values relative to those of Nurr1 deficient brains, suggesting these networks are impaired in Nurr1 deficient brains. (**B, C**) Clusters of significant voxels were reconstructed by voxel-wise group statistics followed by threshold free cluster enhancement (TFCE) to correct for multiple comparisons (see “*Materials and Methods*”). Mean values of independent component (IC) weight in the significant clusters were quantified. The average IC weight in the amygdala network cluster is ≈0.071 (n = 17) in control mice, but significantly reduced to ≈-0.056 (*p* = 0.00090, ***; n = 16) in Nurr1 deficient mice (**B**). The value in the ACC network cluster in control mice is ≈0.41, but reduced to ≈0.030 (by ∼12.51 ± 3.04 folds, *p* = 0.00027, ***) in Nurr1 deficient mice (**C**). **(D-D’ and E-E’)** Seed-based connectivity mapping approach to analyze the specifically connected networks to CLA. The synchronous BOLD signals of amygdala (white arrowheads) and ACC (red arrowheads) networks in Nurr1 deficient brains are weakened in comparison to control brains when either both (**D-D’**) or single (**E-E’**) hemispheric CLA was seeded as regions of interest.

### Nurr1 deficient mice is more resistant to stress-elicited behavioral responses

Neural circuitry between amygdala and CLA contributes to modulating responsive behaviors to stress in rodents^9^. Given the genetic disturbance of CLA and impaired amygdala-CLA communication in Nurr1 deficient mice, we reasoned if Nurr1 was involved in regulation of stress-induced cognitive processing. We first tested control and Nurr1 deficient mice with elevated plus maze (EPM). Nurr1 deficient mice exhibited significantly longer travelling time and distance as well as higher velocity and visit frequency in the open arms of EPM than control mice (Fig. 4A-C, 4F and Fig. S7A). Moreover, Nurr1 deficient mice spent less time in closed arms of EPM than controls despite their similar moving distances (Fig. 4E, 4H). These results implicate Nurr1 deficient mice become less anxious to external stressors. We also defined the open arm ledges (10 cm) as high risk zones^37, 38^, and found that Nurr1 deficient mice were more exploratory as they visited more often the risk zones and travelled longer time and distance in this area than control mice (Fig. 4D, 4G and Fig. S7B), indicating Nurr1 deficient mice are more resistant to the fear of taking risk. We next assessed animals’ anxiety induced by new environment using open field (OF). Nurr1 deficient mice travelled longer time and distance (Fig. 4J, 4K), and exhibited greater locomotion (Fig. S7D) in the central zone of OF than control mice in spite of their statistically indistinguishable velocity and visit frequency (Fig. S7E, S7F), strengthening the conclusions of EPM assay. Collectively, genetic abolishment of CLA ensemble by Nurr1 deficiency makes rodents more immune to stress-elicited emotional responses, such as anxiety and fear.

**Figure 4.**
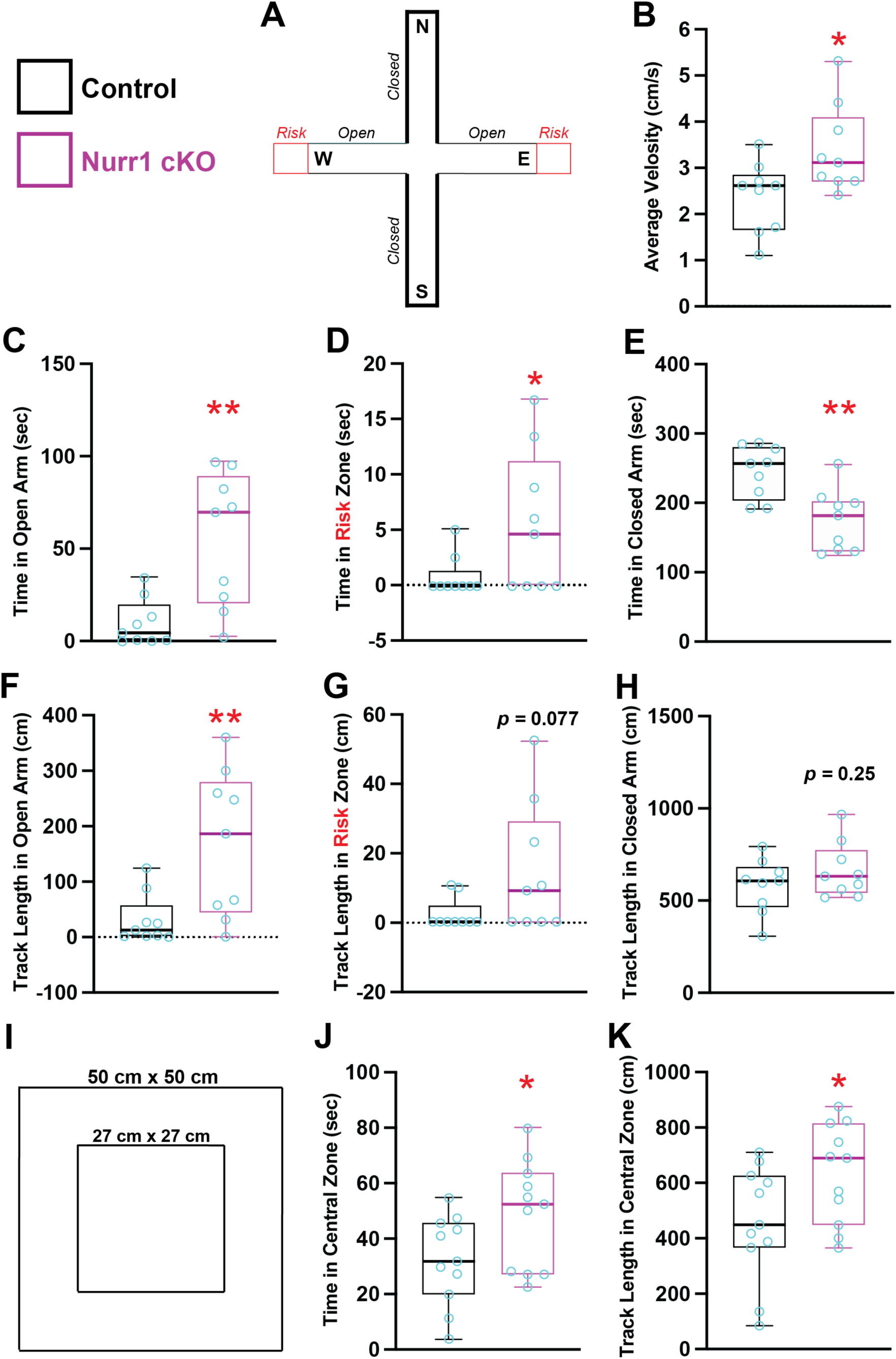
Nurr1 deficient mice are less susceptible when exposing to stressors. (A –. **H)** Analysis of mouse behaviors in elevated plus maze (EPM). (**A**) The apparatus used in EPM (also see “*Methods and Materials*”). The north (N) and south (S) arms are closed with barriers and west (W) and east (E) arms are open arms. The far-end ledge of open arms (10 cm) are defined as risk zones (red). (**B**) The velocity of Nurr1 deficient mice in EPM (≈3.38 cm/s, n = 9) was significantly higher than control mice (≈2.37 cm/s, n = 9), increased by ≈42.71% ± 17.10% (*p* = 0.024, *). **(C – E)** Nurr1 deficient mice spent more time in open arms (≈55.03 sec, n = 9) and risk zones (≈5.56 sec, n = 9) than control mice (open arms: ≈10.18 sec, n = 9; risk zones: ≈0.86 sec, n = 9), increased by ∼4.41 ± 1.25 folds (*p* = 0.0028, **) and by ∼5.49 ± 2.58 folds (*p* = 0.049,*), respectively, whereas Nurr1 deficient mice spent less time in closed arms (≈175.3 sec, n = 9) than control mice (≈244.9 sec, n = 9), decreased by ≈28.41% ± 7.91% (*p* = 0.0024, **). **(F – H)** Accordingly, Nurr1 deficient mice traveled longer distance in open arms (≈167.9 cm) and risk zones (≈14.48 cm) than control mice (open arms: ≈31.42 cm; risk zones: ≈2.28 cm), increased by ∼4.34 ± 1.47 folds (*p* = 0.0095, **) and by ∼5.36 ± 2.83 folds (*p* = 0.077, ns), respectively, but the track length of Nurr1 deficient mice (≈663.9 cm) is comparable to that of control mice (≈579.2 cm) in the closed arms (*p* = 0.25, ns). **(I – K)** Analysis of mouse behaviors in open field (OF) test. (**I**) The schematic view of the open field test apparatus. (**J**) Nurr1 deficient mice spent more time in central zone of OF (≈48.67 sec, n = 11) than control mice (≈32.40 sec, n = 11), significantly increased by ≈ 50.22% ± 23.58% (*p* = 0.046, *). (**K**) The travel distance of Nurr1 deficient mice in the central zone (≈633.4 cm) is longer than that of control mice (≈456.3 cm), increased by ≈38.81% ± 18.12% (p = 0.045, *).

### Transcriptomic profile of claustral neurons regulated by Nurr1

We next aimed to screen downstream effectors of Nurr1 that regulated CLA neuron activities. We isolated the anterior claustro-insular portions of control and Nurr1 deficient cortices (excluding posterior neocortical tissues and amygdala), which were immediately subject to scmRNA-seq. We first generated a heatmap for genes regulated by Nurr1 and found that a wide spectrum of genes altered their expression patterns in Nurr1 deficient mice, particularly in L6-IT cluster (layer 6 intratelencephalon projecting glutamatergic neurons including CLA and dEn cells in Azimuth annotation^39^) (Fig. 5A). ISH for a number of top-rated genes in heatmap validated the specificity of our transcriptomic mapping (Fig. S8). Uniform manifold approximation and projection (UMAP) dimensionality reduction in combination with cell subclass clustering analysis for all cells provided a blueprint of claustro-insular cell heterogeneity, showing that Nurr1 expression in control and Nurr1 deficient tissues was largely overlapping in glutamatergic neuron clusters besides L6-IT cluster (Fig. 5B, 5C). UMAP analysis based on Nurr1+ subpopulations revealed that Nurr1 expression was mainly detected in L2/3 (most likely Arimatsu cells), L6-IT (CLA and dEn) and L6b (Sp) clusters in controls (Fig. 5D, 5E), supported by ISH and cell type specific UAMP (Fig. S1, S8 and S9A). However, a large population of L6-IT cluster cells were no long recognizable in the absence of Nurr1 (red arrowheads in Fig. 5B, 5D), arguing their cell identity shift. Additionally, quite a few CLA-enriched markers, as shown in our data and previous reports^20, 40, 55^, were verified to be downregulated whereas Rorβ was upregulated (Fig. 5F), consistent with our molecular characterization.

**Figure 5.**
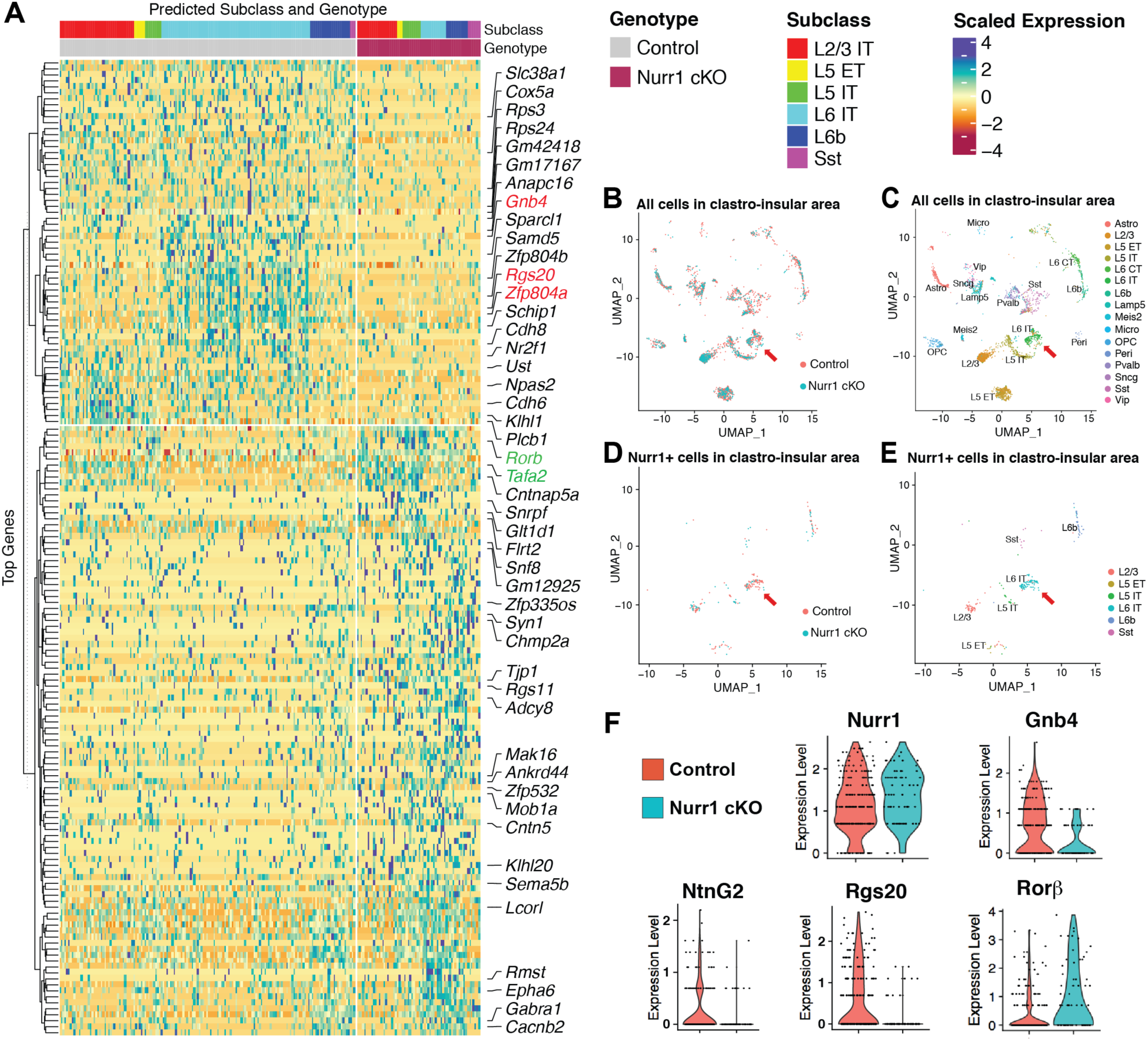
Single cell transcriptomic analysis of Nurr1+ cell populations in claustro-insular region at P0. **(A)** Heatmap of top-rated regulated genes by Nurr1. The map was generated based on single cell transcriptomics of claustro-insular cells in control (left panel) and Nurr1 deficient brains (right panel). Cell subclasses are marked by colors. The figure is separated by expression levels (downregulation: upper panel; upregulation: lower panel) and the expression scales are marked as deep red (-4) as minimal and dark blue (4) as maximal. Some key genes involved in the project are marked (downregulated in red and upregulated in green). **(B, C)** Uniform manifold approximation and projection (UMAP) analysis of the single cell transcriptomes based on all cells isolated from claustro-insular tissues in control and Nurr1 deficient brains (**B**). Each point denotes an individual cell. Claustro-insular cell subclasses in UMAP are grouped and marked by colors in (**C**). Nurr1+ CLA cell population is pointed by a red arrowhead (L6 cluster in Azimuth annotation^39^, the same as **D, E**). Astro, astrocytes; L2/3, layer 2 and 3 cells; L5 ET and IT, layer 5 extra- and intra-telencephalon projecting glutamatergic neurons; L6 CT and IT, layer 6 corticothalamic and intratelencephalon projecting glutamatergic neurons; L6b, layer 6b neurons (Sp); Micro, microglia; OPC, oligodendrocyte precursor cells; Peri, pericytes; GABAergic neuron clusters: Lamp5, Meis2, Pvalb, Sncg, Sst, Vip. **(D, E)** UMAP analysis of Nurr1+ cell populations in claustro-insular region of control and Nurr1 deficient brains. Nurr1+ cell subclasses are marked by colors in (**E**), which are mainly distributed in L2/3, L6 IT and L6b of cortex, consistent with Nurr1 expression pattern. **(F)** Violin plots for selected genes expressed in CLA. The levels of Nurr1-Cterm transcripts in claustro-insular cells of control and Nurr1 deficient brains are equivalent. However, expression levels of Gnb4, NtnG2 and Rgs20 are downregulated in claustro-insular region of Nurr1 deficient brains, whereas the expression level of Rorβ is upregulated.

### Claustral cell positioning relies on suppression of Gαs-PKA signaling by Nurr1

Quite a few numbers of top-rated Nurr1 downstream genes in our transcriptomics were found linked with inhibition of G-protein signaling, such as Gnb4 and Gnb2 (Fig. 2, Fig. 5 and Fig. S9). Additionally, Gng2 (Gγ2), another reduced marker in CLA neurons (Fig. 2B-B’), can heterodimerize with Gnb2/4 (Gβ2/4) to stabilize the silent Gα effectors^41, 42^. We thus hypothesized that low activity of Gα subunits might be vital for CLA development. To test this, we established a gain-of-function strategy by *in utero* electroporation (IUE) to restore target genes in CLA cells at E12.5 (Fig. S10), when the birth peak of mouse CLA neurons is^43^. We first electroporated GFP into control and Nurr1 deficient brains as references and quantified the proportions of Nurr1+/GFP+ cells positioned in CLA area relative to the total double positive cells. While Nurr1+/GFP+ cells were mainly populated in CLA of GFP- electroporated control brains (≈89.94%), only a small fraction of Nurr1-Cterm+/GFP+ cells were still resident in the CLA of GFP-electroporated Nurr1 deficient brains (≈15.44%). Instead, the majority of Nurr1-Cterm+/GFP+ cells in mutant brains were detected in InC area, consistent with their phenotypes in the non-electroporated hemispheres (Fig. 6A, 6B). We then analyzed Nurr1-GFP bicistronically electroporated mutant brains, showing that Nurr1 restoration in Nurr1 deficient CLA cells prevented most of Nurr1+/GFP+ cells from migrating to InC (≈69.63% in CLA) (Fig. 6A, 6B). The fraction should be even larger given some non-CLA cells carrying enforced Nurr1 expression were counted into denominator. Notably, co-electroporation of Gnb4- and Gng2-GFP into Nurr1 deficient brains increased the proportion of Nurr1-Cterm+/GFP+ cells in CLA by ∼2.23 ± 0.24 folds (*p* < 0.0001, ****) relative to GFP-electroporated mutant brains (Fig. 6A, 6B). These results reveal restoration of Gα deactivators in Nurr1 deficient CLA cells facilitates their settlement in CLA territory.

**Figure 6.**
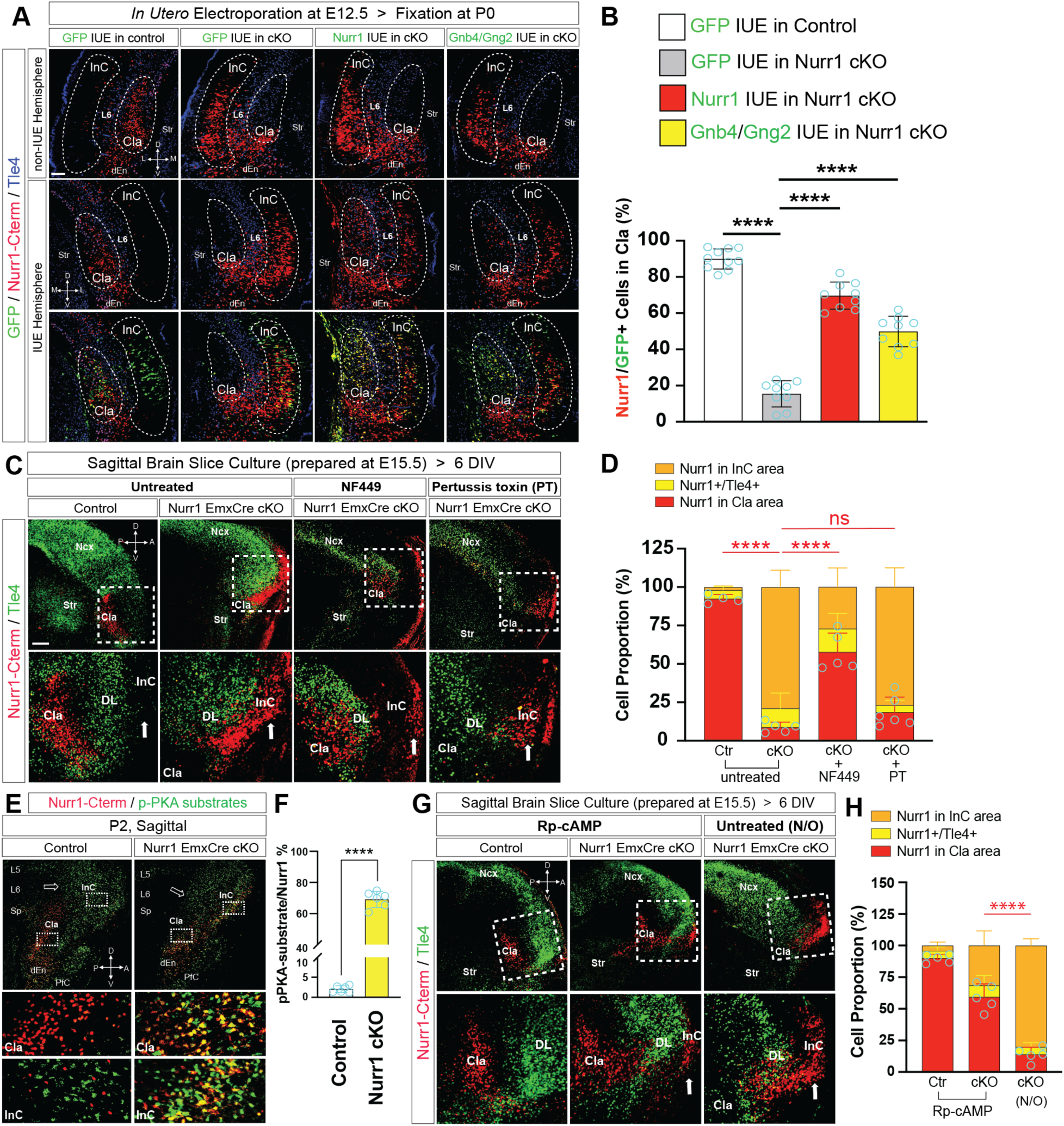
Nurr1 regulates claustral neuron positioning by suppressing Gαs-PKA signaling. (A, **B)** IF for Nurr1-Cterm (red) and Tle4 (blue) on the coronal sections of non-electroporated (non-IUE) and electroporated hemispheres of control and EmxCre mutant brains at P0 (**A**) and quantification for the proportions of Nurr1-Cterm+/GFP+ cells resident in CLA relative to all double positive cells in various electroporation conditions (**B**). The non-electroporated hemispheres serve as comparison references to the electroporated ones of the same brains. IUE of GFP into embryos at E12.5 demonstrates almost all double positive cells reside in normal CLA of control brains (≈89.94%, n = 10), but IUE of GFP into Nurr1 deficient embryos reduces the proportion to ≈15.44% (n = 9), depicting Nurr1 deficient cells largely migrate across Tle4+ L6 cells and disperse in InC. Nurr1-GFP bicistronic electroporation into mutant embryos restricts the Nurr1-Cterm+ cells in the CLA area, elevating the proportion by ∼3.51 ± 0.22 folds (≈69.63% ± 3.47%, *p* < 0.0001, ****; n = 9) relative to that of GFP IUE into mutant embryos. Moreover, Gnb4/Gng2 co-electroporation into mutant embryos restricts a subpopulation of Nurr1-Cterm+ cells in the CLA area, elevating the proportion by ∼2.23 ± 0.24 folds (≈49.89% ± 3.70%, *p* < 0.0001, ****; n = 9) relative to that of GFP IUE into mutant embryos. Notably, the Nurr1-Cterm+ cells in the Nurr1- or Gnb4/Gng2-electroporated mutant hemispheres are visibly less in InC areas than their non-electroporated contralateral hemispheres. The statistical data are analyzed by two-tailed *Student’s t*-test (the same below). Scale bar: 100 µm. **(C, D)** IF for Nurr1-Cterm (red) and Tle4 (green) on the sagittal brain slices of control and EmxCre mutant brains (**C**) and quantification for the proportions of Nurr1-Cterm+ cells resident in CLA (red bars) relative to all Nurr1-expressing cells (**D**). The brain slices were prepared at E15.5 and cultured for 6 days *in vitro* (DIV). The Nurr1+ CLA neurons (≈92.29%, n=4) are properly positioned in control brains, but only a small fraction of Nurr1-Cterm+ cells (≈8.78%, n=5) can be detected posterior to Tle4+ DL neurons in Nurr1 deficient slices. (**D**) The proportion of Nurr1-Cterm+ cells in CLA of mutant brains is increased by ∼5.58 ± 0.65 folds when NF449 (10 μm) was applied (≈57.74% ± 5.72%, *p* < 0.0001, ****; n = 5), but not significantly altered when pertussis toxin (PT, 100 ng/ml) was applied (≈18.65% ± 4.62%, *p* = 0.062, ns; n = 6), relative to that of untreated mutant brains. White arrowheads in (**C**) and (**G**) indicate the Nurr1-expressing cells in InC. Scale bar: 200 µm. **(E, F)** IF for Nurr1-Cterm (red) and phosphorylated PKA substrates (green) on the sagittal sections of control and EmxCre mutant brains at P2 (**E**) and quantification for the proportions of double positive cells relative to all Nurr1-expressing cells (**F**). The Nurr1-Cterm+ cells in Nurr1 deficient brains exhibit tremendously elevated levels of PKA substrate phosphorylation (≈69.07% ± 2.14%, n = 6) compared with that of control brains (≈2.06%, n = 6). **(G, H)** IF for Nurr1-Cterm and Tle4 on the sagittal brain slices and quantification for the proportions of Nurr1-expressing cells in CLA as in (**C, D**). Rp-cAMP (100 μm) treated Nurr1+ cells in control brains are mostly located in CLA (≈90.05%, n=5), and seem normal as that of untreated control brains (≈92.29%, **D**). The proportion of Nurr1-Cterm+ cells in the CLA of Rp-cAMP treated mutant hemispheres is increased up to ≈59.30% ± 5.28% (p < 0.0001, ****; n = 5) in comparison to that of untreated mutant hemispheres (≈13.96%, n = 5).

Gα effector proteins are divided into several subclasses, such as Gαs, Gαi, Gαq and Gα_12/13_, to mediate diverse cell activities, among which homeostasis of Gαs and Gαi cascades primarily regulates cell motility^44^. In order to further study the effect of Gα downstream cascades on CLA cell behaviors, we combined brain slice culture with acute pharmacological treatment. The brain slices were prepared from E15.5 embryos, cultured in supporting media for six days and then fixed by 4% paraformaldehyde^45^. The resulted tissues appeared morphologically intact. Nurr1+ CLA cells in untreated control slices dwelled normally between Tle4+ striatal and InC DL neurons, while the majority of Nurr1-Cterm+ cells in untreated mutant slices migrated across InC DL neurons towards the pia, in agreement with the phenotype by Nurr1 deficiency *in vivo* (Fig. 6C). We then quantified the proportions of Nurr1-expressing cells positioned in CLA or InC relative to the total numbers. There was a huge fraction (≈92.29%) of Nurr1+ cells in the CLA of untreated control slices in contrast to merely ≈8.78% Nurr1-Cterm+ cells in the CLA of untreated Nurr1 deficient slices (Fig. 6D). We next constantly treated Nurr1 deficient slices with NF449, a Gαs-specific cell-permeable inhibitor^46,47^, and found a visibly larger fraction of Nurr1-Cterm+ cells ceasing posterior to InC DL neurons (≈57.74%) (Fig. 6C, 6D). On the other hand, the treatment with pertussis toxin (PT), a Gαi-specific inhibitor^48, 49^, to Nurr1 deficient slices did not exhibit such an effect as NF449 treatment did (Fig. 6C, 6D). Collectively, excessive activity of Gαs signaling in Nurr1 deficient CLA neurons causes, at least in part, their constitutive migration.

PKA signaling, as the most common downstream cascade of Gαs^44, 50^, was assumed to be activated in the context of Nurr1 deficient cells. We indeed found the phosphorylation levels of PKA substrates, but not PKC substrates (Fig. S11A), ectopically surged up in Nurr1 deficient neurons (≈69.13%) in contrast to its barely detectable level in control brains (≈2.06%) (Fig. 6E, 6F). Interestingly, one of catalytic subunits of PKA complex, *Prcacb,* was upregulated in InC (Fig. S11B) but one regulatory subunit, *Prkar2b*, was downregulated in Nurr1 deficient brains (Fig. S9G). To investigate if CLA cell mislocalization in Nurr1 deficient brains is linked with ectopic PKA activities, we prepared brain slice cultures from two hemispheres of the same brain and treated them with or without a cell-permeable PKA inhibitor Rp-cAMP^51^. PKA inhibition by Rp-cAMP enabled a significantly larger population of Nurr1-Cterm+ cells resident in CLA area in the treated mutant hemispheres (≈59.30%) than that of non-treated ones (≈13.96%), mimicking the situation of NF449 treatment. These results suggest minimal activity of Gαs-PKA signaling is crucial for appropriate positioning of normal CLA neurons.

## Discussion

There has been a longtime discussion on if CLA is the main processor for mammalian consciousness, which attracted constantly growing interest in CLA studies in the last two decades. CLA has been confirmed to be the mostly interconnected tissue to coordinate a wide range of sophisticated behaviors as described above. However, molecular mechanisms underlying CLA formation and physiological functions remain little known. Here we report one piece of earliest experimental evidences that Nurr1 is a master regulator to orchestrate CLA morphogenesis, cell identity determination, FC networks and cognitive behaviors.

The evo-devo relation between CLA and InC, and CLA cell migration path were not in agreement until Charles Watson and Luis Puelles proposed the latest descriptive model based on Nurr1 expression pattern, hypothesizing that principal CLA neurons derived from lateral pallial progenitors (LPP) first populate ventrally to Sp and later arriving cells bypass CLA to form InC in a typical inside-out manner^1, 19^, but direct experimental evidence for this model still lacks. Here we confirm this theory using our mouse model where CLA cells deficient for Nurr1 transcription regulation constitutively migrate radially along radial glia into InC and are specified into InC-like cells. These results suggest CLA and InC derive from the same lineage and migrate along the same radial columns, and that Nurr1 is the decisive factor to segregate the earlier coming clusters (mainly at E12.5)^43^ to form subcortical CLA from the later arriving InC neurons. Nevertheless, we did not observe any tangentially migrating Nurr1-Cterm+ cells in mutant brains throughout our study, challenging their point of view that some Nurr1+ cells may migrate dorsally to become Sp or Arimatsu cells. In contrast, Nurr1 deficient CLA cells constitutively migrated along a radial path and disconnected with intact Sp, raising a myth if CLA is truly a spatially continuous structure of Sp in spite of their partially common evolutionary origin^20^.

Normal CLA territory has been described to be surrounded by striatum and Tle4+ InC DL neurons in mouse brains where a clear extreme capsule structure lacks^52^. CLA cells in Nurr1 deficient mice easily cross the barrier to invade InC whereas surrounding DL cells appear to occupy original CLA territory. These results suggest there is mutually exclusive interaction between CLA and InC cells to assist claustro-insular organization. Nurr1 presides over CLA party and its deficiency alone can disrupt claustro-insular cytoarchitecture. Notably, our scmRNA-seq data revealed quite a few cell-cell contact clues that were dysregulated by Nurr1 deficiency, such as molecules in ephrin-Eph and semaphorin-plexin signaling. These proteins are often axon membrane bound to mediate *trans* repelling interplay between contacted tissues^53, 54^. This may provide a molecular hint for how CLA is delineated by white matter (external and extreme capsules) in human brains.

Additionally, we have found Nurr1 not only regulates CLA morphogenesis but also its cell fate specification, by showing that CLA signature genes are selectively downregulated while InC-enriched genes, such as Cyp26b1, Rorβ and Neurod1, are significantly upregulated in Nurr1 deficient CLA cells. Consistently, single cell transcriptomic analysis verifies our molecular characterization, and further reveals that CLA specific genetic program is broadly affected by Nurr1 deficiency. Moreover, another group also reported similar transcriptomic results when we were preparing the manuscript^55^. These independent studies thus confirmed the same conclusion that Nurr1 is a key player to program CLA identity. Remarkably, not all cells carrying enforced Nurr1 expression by IUE were converted into CLA neurons, suggesting that Nurr1 alone is necessary but may not be sufficient to determine CLA cell identity. Nurr1 is also strongly expressed in Sp and Sb, but its absence does not affect their cell identity or anatomic structure. Nurr1’s role in postmitotic control of cell fate is plausibly cell type- or region-specific.

Nurr1 expression has been detected in both cortical progenitors and CLA neurons in early brain development (*Allen brain atlas*), posing a question if Nurr1 governs CLA cell activities pre- or postmitotically. We answered this question by genetic depletion of Nurr1 at distinct stages. Nurr1 inactivation in postmitotic neurons (Nex-Cre) could fully recapitulate the CLA developmental phenotypes as those when Nurr1 was inactivated in progenitors (Emx-Cre), indicating a postmitotic mechanism of Nurr1 in the context of CLA formation. Moreover, IUE of GFP into wild type brains (Fig. S10) showed that Nurr1 expression was not present in CLA cells during migration from VZ to CP, suggesting that Nurr1 expression is separated into two stages during CLA development – progenitor and neuronal, and that Nurr1 expression in progenitors is dispensable for cell migration towards CLA but its later expression in neuronal stage is the key for CLA morphogenesis.

Nurr1 is not only critical for CLA development, also for CLA-linked inter-areal connectivity organization. For instance, fMRI analysis seeding CLA as ROI showed synchronized activity of amygdala network was impaired in Nurr1 deficient mice. Recent report showed that stress responsive behaviors controlled by amygdala relied on CLA ensemble as part of processing circuit^9^. To test the idea if normal CLA integrity and FC regulated by Nurr1 is required for CLA dependent behaviors, we assessed our mice by EPM and OF assays. Nurr1 deficient mice were clearly more insensitive to stress-elicited emotional responses, consistent with their argument that CLA neuron inactivation led to increased resilience to stressors^9^. Taken together, Nurr1 is seated high in genetic hierarchy in CLA to dictate its entire development-connectivity-cognition dimension, in line with its life-long expression in CLA.

Immense evidence has shown G-protein mediated signaling is one of major regulatory pathways in cell motility, but Gα subclass effectors play complex and contextually divergent roles in variable cell types^56^. Both Gαs and Gαi isoforms have been shown to stimulate promigratory events in certain cell types but suppress cell migration in other circumstances, depending on their expression levels, homeostasis of Gαs/Gαi, and cell type-specific niche^56–61^. We therefore utilized selective inhibitors to reveal Gαs-PKA axis to be involved in the context of CLA cell aberrant migration caused by Nurr1 deficiency. Intracellular PKA pathway, as a main responsive signaling of Gαs, serves as a central paradigm to regulate cytoskeleton dynamics so as to coordinate almost all key events of cell migration, such as polarization, adhesive interaction with extracellular matrix, leading edge formation and soma translocation^62, 63^. For instance, Gαs-PKA signaling facilitates embryonic cortical neuron migration by directly phosphorylating the microtubule-associated protein *Doublecortin (Dcx)* to induce promigratory cytoskeleton reorganization^64^. Despite the delicate spatio-temporal control of PKA signaling in a motile cell in terms of its abundance, equilibrium of catalytic and regulatory subunits, activation timing and subcellular compartmentation, basic PKA activity is generally required^62–66^. In this regard, to simply minimize intracellular PKA activity may be the most cost-efficient implementation to ensure cell motionless. Nurr1 sets serial barriers at transcription level to keep PKA activity beneath threshold, such as upregulating upstream repressors and PKA regulatory subunits but downregulating catalytic subunits. Consistently, PKA substrate phosphorylation was strongly detected in radially migrating neocortical neurons, but selectively absent in static CLA cells.

Given the deep position and tight interplay with neighboring tissues of CLA (and dEn) in mouse brains, cell type-specific Cre line logically serves as a better tool for genetic study than artificial lesions^67^, which debilitates complete and neat gene inactivation in CLA cells. However, the CLA-specific Cre lines that have been reported in an experiment so far trigger recombination either in a relatively small subset of CLA neurons (Egr2-Cre) or only in adult CLA (Tbx21-Cre)^4, 10, 68^. Our study would open a new window for cell type-specific investigation of CLA (and dEn) in the future.

## Acknowledgments

We want to thank Rike Dannenberg, Denis Lajkó and Jutta Schüller for their brilliant technical support. NEX-Cre mice were kindly contributed by the group of KA Nave, Max-Planck-Institute of Experimental Medicine, Göttingen, Germany. Joseph Kuchling provided necessary assistance to upgrade the fMRI pipeline for seed-back analysis. We also want to thank Markus Aswendt for sharing the standard operating procedures for rs-fMRI anesthesia.

## Author contributions

Conceptualization: KY, VT

Methodology: KY established the cellular and surgical methods, such as brain slice culture, IUE, et al. KY and PL developed some of the molecular and cellular methods, such as ISH, IF and DiI, et al. AN analyzed the transcriptomic data. SM and MF performed fMRI examination. MM, CF and PBS introduced and upgraded the analytic pipeline of fMRI. SPK and PBS analyzed the fMRI data. KY, PD and ML developed the animal behaviors tests. KY, PL and PD performed genotyping.

Investigation: KY and PL performed molecular and cellular characterization for Nurr1 cKO mouse model. KY carried out IUE and brains slice culture experiments. KY isolated the cortical tissues and AN analyzed the scmRNA sequencing data. SM, MF, SPK and PBS contributed to the experimental line of fMRI. KY and PD performed animal behaviors tests under ML’s supervision and KY analyzed the statistical data.

Funding acquisition: 1) Main funding of the project for molecular characterization of mouse model, sequencing and animal behavior tests was provided by VT; 2) funding for fMRI analysis was provided by PBS.

Supervision: KY, VT

Writing original draft and editing: KY, VT

## Data availability

All data in the main text and supplement supporting the scientific findings of this study will be made available on a central repository.

