## Supplemental Materials for "Nurr1 orchestrates claustrum development and functionality"

### Supplementary Materials

#### Materials and Methods

##### Mice

Nurr1$^{flox/flox}$; EmxCre$^{cre/wt}$ and Nurr1$^{flox/flox}$; NexCre$^{cre/wt}$ mice were obtained by crossing Nurr1$^{flox/flox}$ with Emx-Cre and Nex-Cre, respectively. The control mice used in this study were Cre-negative Nurr1$^{flox/flox}$ mice (Nurr1$^{flox/flox}$; EmxCre$^{wt/wt}$ or Nurr1$^{flox/flox}$; NexCre$^{wt/wt}$ mice). The genotyping strategies for these lines were reported previously[1,2] and the genotyping primers are listed in the Tab. S1. The term 'Nurr1 deficient mice' in text means Nurr1 conditionally deficient mice recombined by EmxCre (Nurr1$^{flox/flox}$; EmxCre$^{cre/wt}$) unless noted otherwise. Experiments involving living mice conform to German regulatory standards and were approved by the authority (LaGeSo Berlin).

##### Molecular cloning and constructs

The plasmid backbone used in this study is a Cre-dependent bicistronic expression vector (pCAG-FPF-GFP) as previously described[3]. The full-length open reading frames of Nurr1, Gnb4 and Gng2 with their original kozak sequences were subcloned into pCAG-FPF-GFP plasmids by proper restriction enzymes. The constitutive expression vector pCAGIG-GFP was used for wild type mice (fig. S10). All cloned genes in this study were verified by sequencing. Cloning primers are listed in Tab. S1.

##### Immunofluorescence

The mouse brains for immunofluorescence (IF) were fixed in 4% paraformaldehyde (PFA, diluted in DEPC-treated PBS, DPBS) over night (O/N) and subsequently incubated in DPBS containing 15% (for 6 hrs) and 25% sucrose O/N at 4°C. The fixed brains were embedded in Tissue-Tek OCT and sectioned at -20°C (16 μm in thickness). The brain sections were incubated in blocking solution (BS: 10% horse serum, 2% BSA, 0.5% Triton X-100 in PBS) for 1 hour (hr) following incubation with primary antibodies (in BS) O/N at 4°C. On the second day, the sections were incubated with secondary antibodies for 1 hr. The sections were washed 3 times in PBS between each incubation step (10 min per wash). Primary antibodies used in IF are listed in Tab. S2. Fluorescent secondary antibodies used in IF are all from Jackson ImmunoResearch (all were raised in donkey and diluted 1:500).

**Nissl staining**

The embedded brains were cut into sagittal or coronal sections at thickness of 16 µm and rehydrated in 1x PBS for 15 min. The sections were stained in 0.5% cresyl violet solution for 15 min at RT and washed three times with 1x PBS. Eventually, the stained sections were dehydrated in an ascending alcohol series (50%-80%-90%-100%, 5 min each) and 100% Xylol for 5 min, and then mounted in Permount toluene media.

***In situ* hybridization**

The mouse brains for *in situ* hybridization (ISH) were processed as in IF. First Day: brain sections were dried in vacuum for 30 min, fixed in 4% PFA (in DPBS) for 15 min and incubated in proteinase K solution (20 mM Tris pH 7.5, 1 mM EDTA pH8.0, 20 µg/ml proteinase K) for 2.5 min. Subsequently, brain sections were washed in 0.2% Glycine (in DPBS), post-fixed in 4% PFA containing 0.2% glutaraldehyde (SIGMA) for 15 min, and followed by prehybridized in hybridization buffer (HB: 50% deionized formamide, 5x SSC, 1% blocking reagent from Roche, 5 mM EDTA, 0.1% Tween20, 0.1% CHAPS, 100 µg/ml Heparin, 100 µg/ml yeast RNA, 50 µg/ml Salmon sperm DNA) at 65°C for 2 hrs. Eventually the slides carrying targeted probes (in HB) were incubated at 68°C O/N. The sections were washed twice in DPBS between each step (5 min per wash).

Second Day: the brain sections were washed once in 2x SSC, incubated in RNase solution (0.5 M NaCl, 10 mM Tris pH 8.0, 20 µg/ml RNase A) for 30 min at 37°C, washed once in 2x SSC, and then washed stringently 3 times in 50% formamide/ 2x SSC at 63°C (30 min each), and eventually washed 3 times in KTBT buffer (50 mM Tris pH7.5, 150 mM NaCl, 10 mM KCl, 1% Triton X-100) for 10 min each. Subsequently, the sections were incubated in blocking solution (BS, KTBT containing 20% sheep serum) for 2 hrs, followed by incubation with anti-digoxigenin antibody (conjugated with alkaline phosphatase, 1:1500) in BS at 4°C O/N.

Third day: the brain sections were washed 3 times in KTBT for 30 min each, washed twice in NTMT buffer (100 mM Tris pH 9.5, 100 mM NaCl, 50 mM MgCl2, 0.1% Tween 20) for 15 min each, and were eventually incubated with NBT/BCIP substrates (in NTMT). The staining was monitored hourly until the signals showed up. The stained sections were subjected to an ascending alcohol series (50%-80%-90%-100%, 5 min each), clearing solution (benzyl alcohol : benzyl benzoate = 1 : 2) for 15 min, and finally mounted in Permount medium.

**Retrograde DiI labeling**

The mouse brains (P5 or P12) were fixed in 4% PFA at 4°C O/N, followed with twice washes with 1x PBS. The lipophilic DiI crystal was carefully buried in the primary motor cortex (P12, intact brains) and hypothalamus (P5, remove about 1/5 of caudal portions of brains in length to expose the hypothalamus) of fixed brains, which were subsequently incubated in 1x PBS containing 0.2% sodium azide ($NaN_3$) in the dark at 37°C for 3 weeks. The brains carrying DiI were sectioned at 200 μm in thickness by a vibratome, which were then subject to IF.

***In utero* electroporation**

*In utero* electroporation (IUE) was performed as previously described[3]. During IUE, the pregnant mice carrying E12.5 embryos were kept laid down on a heating pad and anesthetized by constant inhalation of isoflurane blended with oxygen. Subcutaneous administration of tamgesic was done before an operation was started. The embryos were gently pulled out with ring-headed forceps from an incision (≈15 mm) along the abdomen midline. DNA constructs (500 ng/μl each, mixed with fast green dye) in a glass capillary was enforced by vacuum pico-pump into either side of cerebral lateral ventricles. The electrodes were positioned ventro-caudally to biauricular in order to target claustrum (anode at the side of injection). Electroporation was achieved by an electroporator (setup: 6 times pulses, 30V voltage, 40 ms pulse duration and 999 ms interval time). PBS containing antibiotics (100 units/ml Penicillin-Streptomycin) was applied to each operated embryo right after electroporation. After surgery, the mice were put back to an individual cage marked with electroporated genes and dates. The operated mice were monitored every day until sacrifice at P0.

**Organotypic brain slice culture**

The protocol was modified from previously described[4]. The embryonic brains at E15.5 were dissected out and separated into two hemispheres from the midline, which were placed on the sagittal plane and immediately embedded in low melting-point agarose (kept at 45°C before use) in a plastic mold. The brains were sliced in cold Krebs buffer (12.6 mM NaCl, 0.25 mM KCl, 0.12 mM $NaH_2PO_4$, 0.21 mM $CaCl_2$, 0.12 mM $MgCl_2$, 100 mM HEPES buffer, 0.5 mg/ml Gentamicin, 100 units/ml Penicillin-Streptomycin) using a vibratome (300 μm in thickness). The brain slices were gently transferred by blunt-ended microspatulas onto a Millicell cell culture insert (PET membrane, pore size 1 μm) in a well of 6-well plate and incubated in 1m MEM medium (10% fetal bovine serum, 0.5% Glucose,100 units/ml Penicillin-Streptomycin in MEM solution) at the bottom for 1 hr in a cell culture incubator (37°C and 5% $CO_2$). Subsequently, the slices were cultured in Neurobasal medium (5% fetal bovine serum, 2% B27 supplement, 1% N2 supplement, 0.5% Glucose, 1% Glutamine, 100 units/ml Penicillin-Streptomycin in Neurobasal solution) with or without inhibitors in the incubator for 6 days. The medium was renewed every 24 hrs. The brain slices were fixed by 4% PFA O/N and washed in PBS 3 times (1hr each) after culture. The tail of each embryo was collected for genotyping.

**Functional magnetic resonance imaging**

1) *Experimental procedures*: anesthesia was induced by 2% isoflurane in a mixture of 80% air and 20% O2. A subcutaneous catheter was placed and freely-breathing animals were measured at 7 T (BioSpec 70/20 USR, Bruker, Germany) with a Tx/Rx 1H-cryoprobe and ParaVision 6.0.1 software. The protocol consisted of localizer scans, B0 mapping and shimming, T2w MRI, and resting state functional magnetic resonance imaging (rs-fMRI). A bolus of medetomidine (0.1 mg/kg subcutaneously, Cepetor, CP Pharma) was applied during B0 mapping and isoflurane gradually reduced to 0.5% over 3-5 min and kept at least for 5 min at 0.5% isoflurane before the start of rs-fMRI adapting the protocol as previously reported[5]. During the switch of anesthesia regimes, T2w images were acquired with a 2D-RARE sequence with repetition time/echo time (TR/TE) = 4300 ms / 33 ms, RARE factor 8, 2 averages, 40 contiguous axial slices with a slice thickness of 0.4 mm, field of view (FOV)=19.2x19.2 mm2, matrix (MTX)=192x192, bandwidth (BW)=34.7 kHz and total acquisition time (TA)=3:26 min. Rs-fMRI images were acquired with a 2D gradient echo EPI sequence (TR/TE = 1000 ms / 13 ms, flip angle FA=50°, 16 contiguous axial slices with a slice thickness of 0.75 mm, FOV=19.2 x 12 $mm^2$, matrix of 128x80, BW=400 kHz, 300 repetitions and TA=5:00 min).

2) *MRI data analysis*: Rs-fMRI data were preprocessed in RABIES (https://github.com/CoBrALab/RABIES)[6]. Briefly, data were corrected for motion, susceptibility distortion and signal that was confound-corrected via signal confound regression taking the signal from cerebral spinal fluid. The cleaned time courses were then processed using two different approaches. In the first approach, analysis within each functional network was carried out using the dual-regression framework as described previously in the context of mice[7, 8]. Briefly, spatially delineated non-thresholded reference maps were used to extract a weighted average BOLD time course for each scan. A general linear model was used to regress these time courses into the individual scans, to obtain parameter estimate maps indicative of network strength at every voxel for each corresponding reference map. The spatial reference maps used to delineate networks to be tested in the dual-regression analyses were 17 reference resting state networks (including piriform cortex, sensory cortex, motor cortex, barrel cortex,

limbic cortex, visual cortex, auditory cortex, anterior cingulate cortex, retrosplenial cortex, hippocampus, striatum, amygdala, thalamus, et al.) as determined in a previous study with group independent component analysis (ICA)[9]. Voxel-wise statistics on the parameter estimate maps from dual regression were performed using permutation test and corrected for multiple comparison using the threshold free cluster enhancement (TFCE) approach. For the analysis of connectivity changes in specific regions of interest (ROI), the mean signals of ROI were correlated with all other voxels in the brain in a seed-based approach for each animal. Since the DSURQE mouse atlas underlying RABIES did not contain all ROI, a brain atlas was derived from the Allen brain atlas (CCFv3) containing a custom set of regions (such as claustrum, insular cortex, anterior cingulate cortex, orbital cortex, amygdala, prelimbic cortex, retrosplenial cortex, hippocampus, motor cortex, visual cortex, auditory cortex, et al.) and was registered to the RABIES template space using ANTx2 (https://github.com/ChariteExpMri/antx2)[10]. Voxel-wise group statistics on the seed-based connectivity maps was performed using permutation test and corrected for multiple comparison using the TFCE approach.

**Single cell mRNA sequencing**

Briefly, the claustro-insular portions of isolated mouse cerebral cortices (amygdala, hippocampal and posterior cortical tissues were manually excluded) were dissociated using ‘Neural Tissue Dissociation Kit (P)’ (Miltenyi Biotec) containing 1x protease inhibitors (SIGMA) according to the manual instruction. The cell suspension was then incubated in ‘nuclei EZ lysis buffer’ (SIGMA) for 5 min on ice and quickly passed through single cell strainers (pore size 70 µm, Miltenyi). Approximately 10,000 nuclei were procured for single-nuclei encapsulation using a 10X single cell controller. 10x Genomics 3' v3 gene expression (GEX) libraries were prepared and sequenced on a single Novaseq sequencing system (S1 lane). Transcript reads were aligned to GRCm38 (mm10) genome reference consortium using cell ranger, and the reads were analyzed using Seurat quantification pipeline in combination with Azimuth annotation[11].

**Animal behavior analysis**

1) *Open field*: the open field test is used to assess general locomotor activity levels and anxiety in rodents. On the day of testing, the animals were moved to a neighboring holding room 30 min before the test in order to allow animals to acclimate to the room conditions. Temperature was maintained between 20-24°C, humidity between 45-65%, ventilation, noise intensity and lighting intensity was also kept at appropriate levels for mice for the duration of the experiments (the same to elevated plus maze). The maze size is 50 cm X 50 cm and the center size is 27 cm X 27 cm. The mouse was gently

placed at the peripheral area when a test begun. The program was set as delay time (10 sec) followed with testing time (10 min). All its movement was recorded by a camera (video encoder) with “tracking mode” of the “Viewer” software (BIOBSERVE), which also carried out face-tail validity, head direction and body centroid (the same to elevated plus maze). The amount of time spent in central zone of the arena, travel distance in the central and peripheral zones, number of visits to the center and average velocity are measured. The apparatus was cleaned by 5% EtOH after each test. The test does not discriminate sexes.

2) *Elevated plus maze*: this test is used to assess anxiety-related responses in rodents. The maze apparatus consists of 2 open arms and 2 closed arms, of which each is 50 cm in length and white in color. The barriers of closed arms are 15 cm in height. The far-end 15 cm of open arms is defined as risk area. The maze is elevated on a 40 cm-high platform. A mouse was gently placed at the joint area of open and closed arms (central area) in the beginning of a test and allowed to explore for 5 min. The mouse behaviors were also recorded the “Viewer” software. The amount of time spent in open and closed arms as well as in rick zones, the travel distance in each area, the number of visits to each area and average velocity were measured. The time and track length in risk zones were exclusive to those in open arms. The apparatus was cleaned by 5% EtOH after each test. The test does not discriminate sexes.

**Quantification and statistics**

Quantification between control and Nurr1 deficient groups was analyzed with two-tailed *Student's t*-test. The charts present mean values ± standard deviation (SD). Significance: p<0.0001, ****; p<0.001, ***; p<0.01, **; p<0.05, *; p>=0.05, ns.

#### Supplement figures

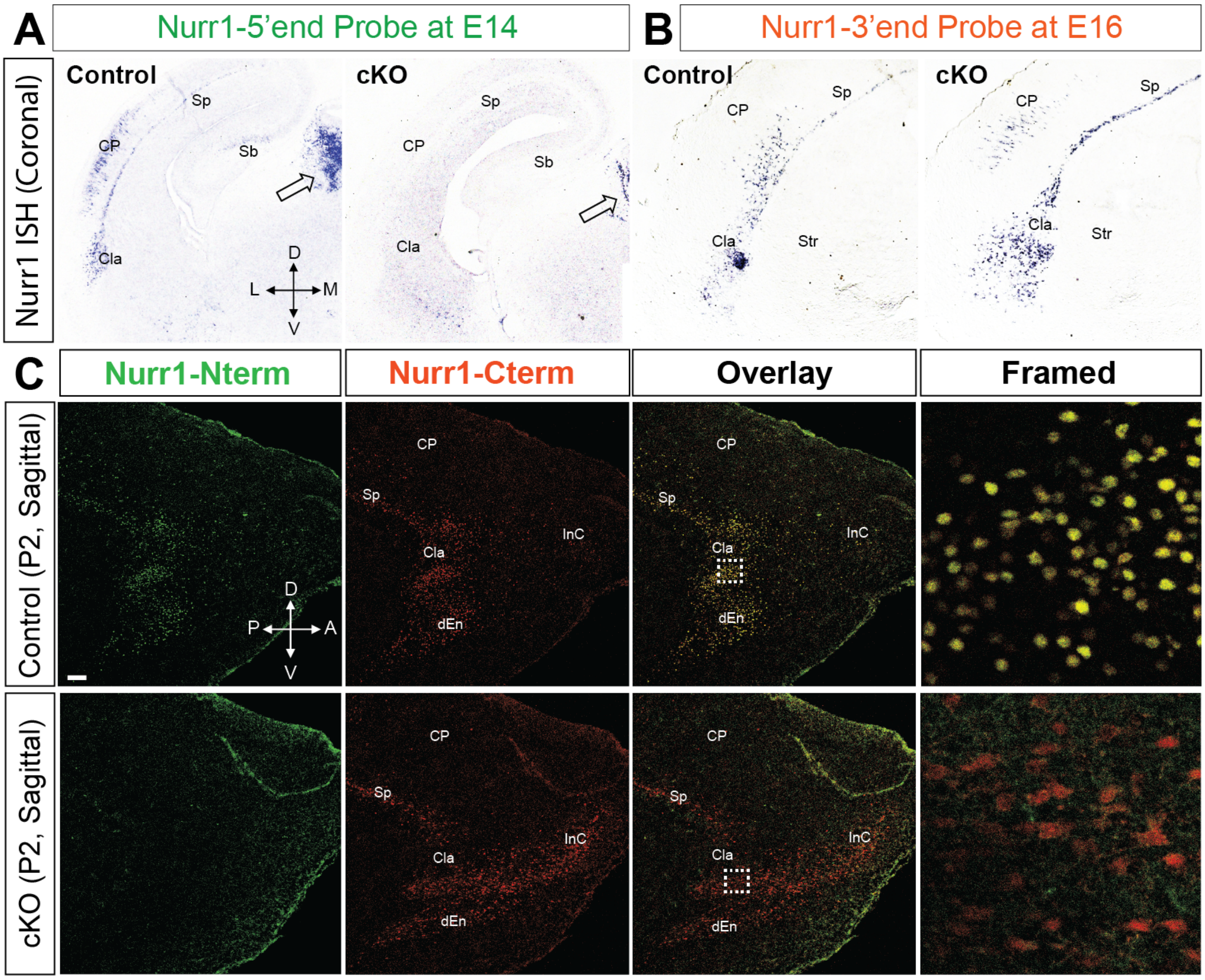


**Figure S1 The C-terminal truncate of Nurr1 is traceable by antibody and *in situ* hybridization probe in Nurr1 transcription-deficient cells.**

**(A, B)** *In situ* hybridization (ISH) using the Nurr1-5'end probe (**A**, at E14) or Nurr1-3'end probe (**B**, at E16) on coronal sections of control and EmxCre+ Nurr1 conditional deficient (cKO) brains shows Nurr1 is expressed in the developing subiculum (Sb), subplate (Sp), claustrum (CLA) and cortical plate (CP) in control mice, but ISH signal is undetectable in the cortex of Nurr1 deficient brains (**A**). The arrowheads indicate ISH signals in the habenula of control and Nurr1 deficient brains as internal reference. **(B)** The ISH signals of Nurr1-3'end probe at E16 are detectable in both control and mutant brains. Str, striatum. D, dorsal; M, medial; V, ventral; L, lateral.

**(C)** Immunofluorescence (IF) on sagittal sections of control and Nurr1 deficient brains at P2 shows that the Nurr1-Nterm (green) and Nurr1-Cterm (red) antibodies (ab) label the neurons in CLA, dorsal endopiriform nucleus (dEn) and Sp in control mice. The Nurr1-Nterm ab shows negative signal in mutant brains, however, Nurr1-Cterm ab remains detectable signals in CLA, dEn and Sp, consistent with ISH. The overlay images show that all Nurr1-Cterm+ CLA cells are double positive with Nurr1-Nterm ab signals (yellow) in controls, suggesting the high specificity of both ab. Nurr1-Cterm polypeptide displays cytoplasmic localization due to loss of its nuclear localization signal (aa 287 – 314). The framed images show magnification views of the dotted-line boxed areas (the same below). InC, insular cortex. A, anterior; P, posterior. Scale bar: 200 µm.

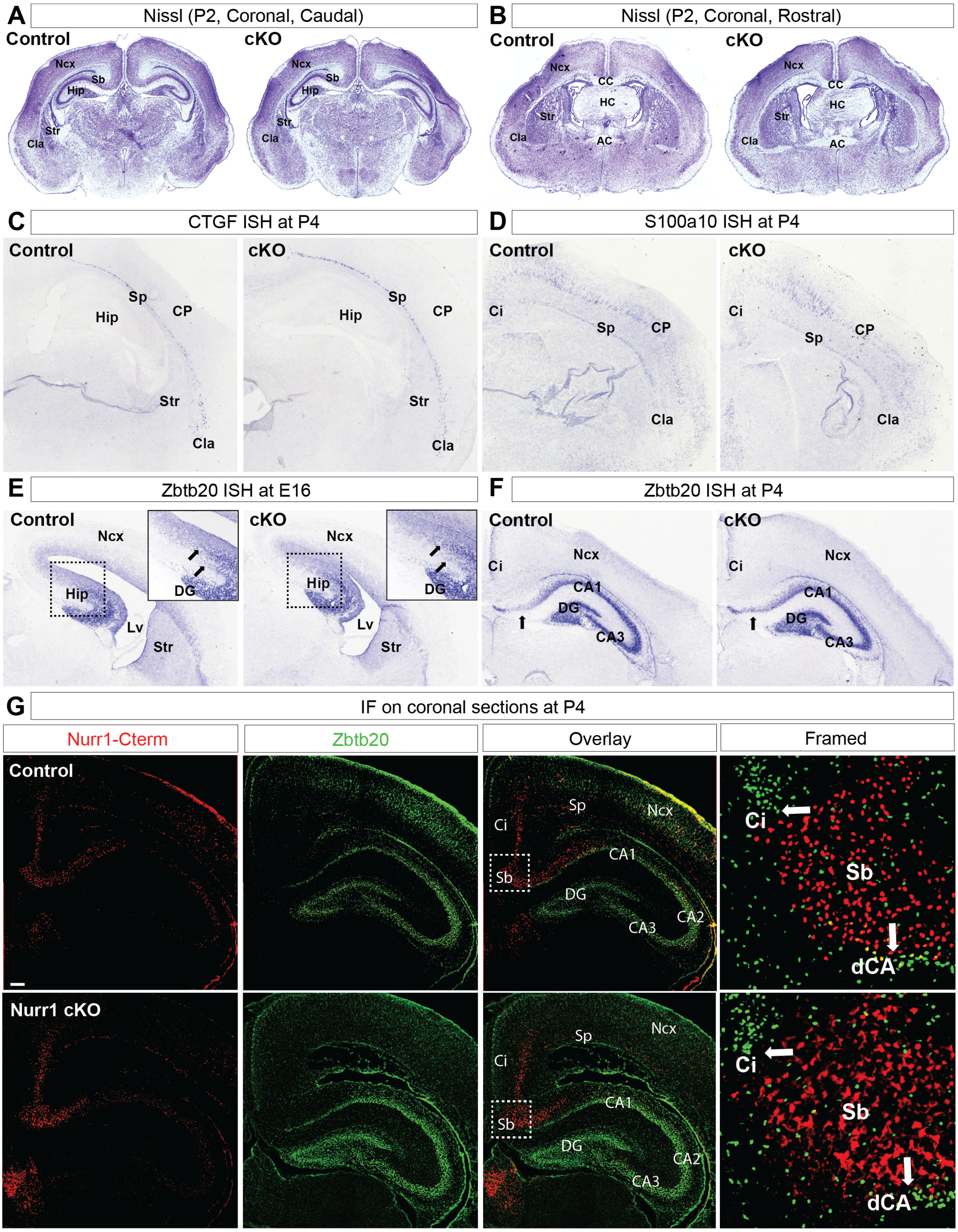


**Figure S2 Nurr1 deficiency does not affect morphology of subplate and subiculum.**

**(A, B)** Nissl staining on caudal (**A**) and rostral (**B**) coronal sections of control and Nurr1 deficient brains shows that the main cerebral structures of Nurr1 deficient mice are basically normal. Hip, hippocampus; Ncx, neocortex; AC, anterior commissure; CC, corpus callosum.

**(C, D)** ISH using the CTGF (**C**) and S100a10 (**D**) probes on coronal sections of control and Nurr1 deficient brains at P4 shows that the Sp is not disrupted in mutant brains. Ci, cingulate cortex.
**(E, F)** ISH using the Zbtb20 probe on coronal sections of control and Nurr1 deficient brains at E16 (**E**) and P4 (**F**) shows that the borders between Ncx and Hip are unaltered in Nurr1 deficient brains. The arrowheads in (**E, F**) indicate the neocortico-hippocampal borders. Lv, lateral ventricle; DG, dentate gyrus; CA1 and CA3 indicate the *Ammon's* horn subareas.

**(G)** IF staining for Nurr1-Cterm and Zbtb20 on coronal sections of control and Nurr1 deficient brains at P4 shows that Nurr1 expression in Sb separates hippocampal from neocortical structures in both control and Nurr1 deficient brains. The framed images show magnification views of the boxed areas in overlay. The arrowheads indicate the neocortico-hippocampal borders. dCA, distal area of *Ammon's* horn CA1. Scale bar: 200 µm.

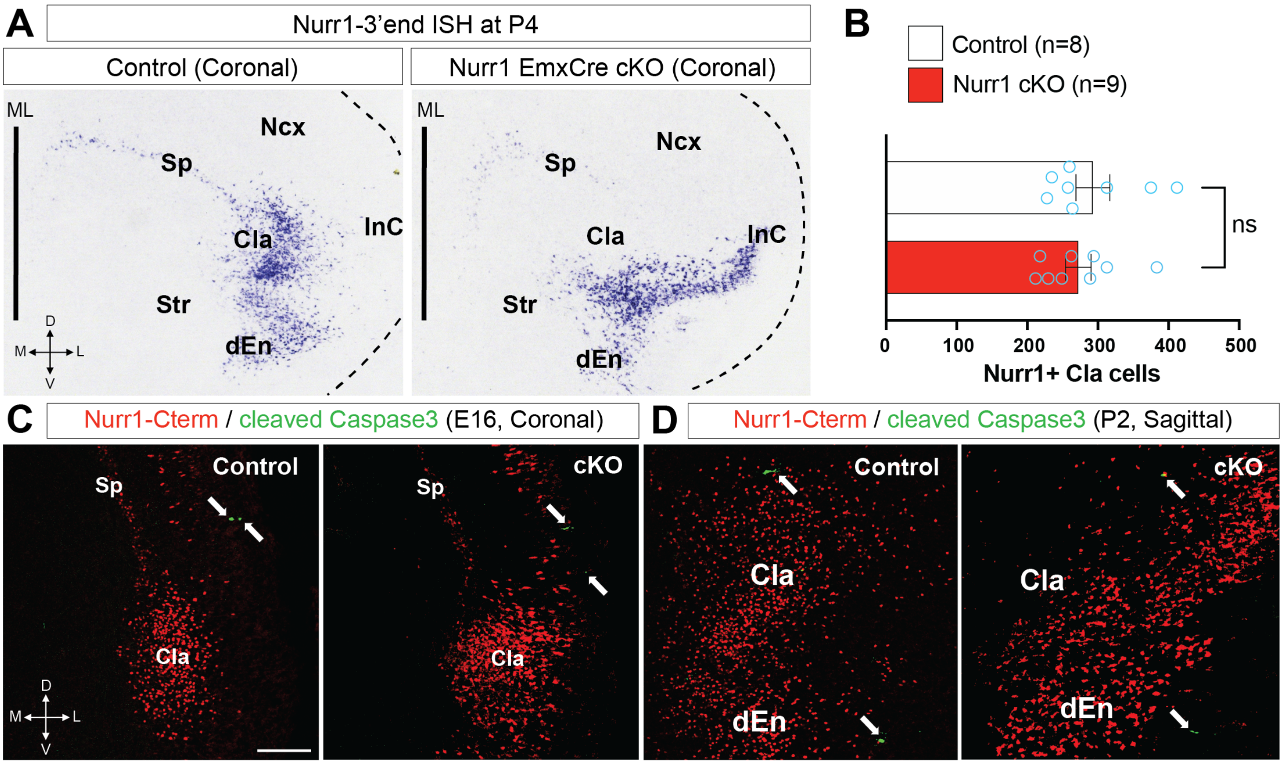


**Figure S3 Nurr1 does not affect claustral cell production or survival.**

**(A)** ISH using the Nurr1-3'end probe on coronal sections of control and Nurr1 deficient brains at P4. Nurr1+ CLA cells form a compact crescent-like structure in control brains but these cells over-migrate in InC and thus transformed into a wing-like structure in Nurr1 deficient brains. ML, mid-line. The dotted lines delineate the tissue borders.

**(B)** Quantification for Nurr1+ CLA neurons based on Nurr1 IF on coronal sections of control and Nurr1 deficient brains at P2. Each value is the average of two sections from the same brain. The mean value of Nurr1+ CLA cells in control brains is ≈293.0 (n = 8), while the mean value of claustro-insular Nurr1-Cterm+ cells in mutant brains is ≈272.2 (n = 9). There is no significant difference between control and mutant brains by two-tailed *Student's t*-test ($p = 0.50$, ns).

**(C, D)** IF for Nurr1-Cterm (red) and cleaved Caspase3 (green) on E16 coronal sections and P2 sagittal sections of control and Nurr1 deficient brains shows only a few apoptotic cells in either control or mutant brains, suggesting that Nurr1 does not affect survival/apoptosis of CLA cells. The arrowheads indicate cleaved Caspase3 positive signals. Scale bar: 200 µm.

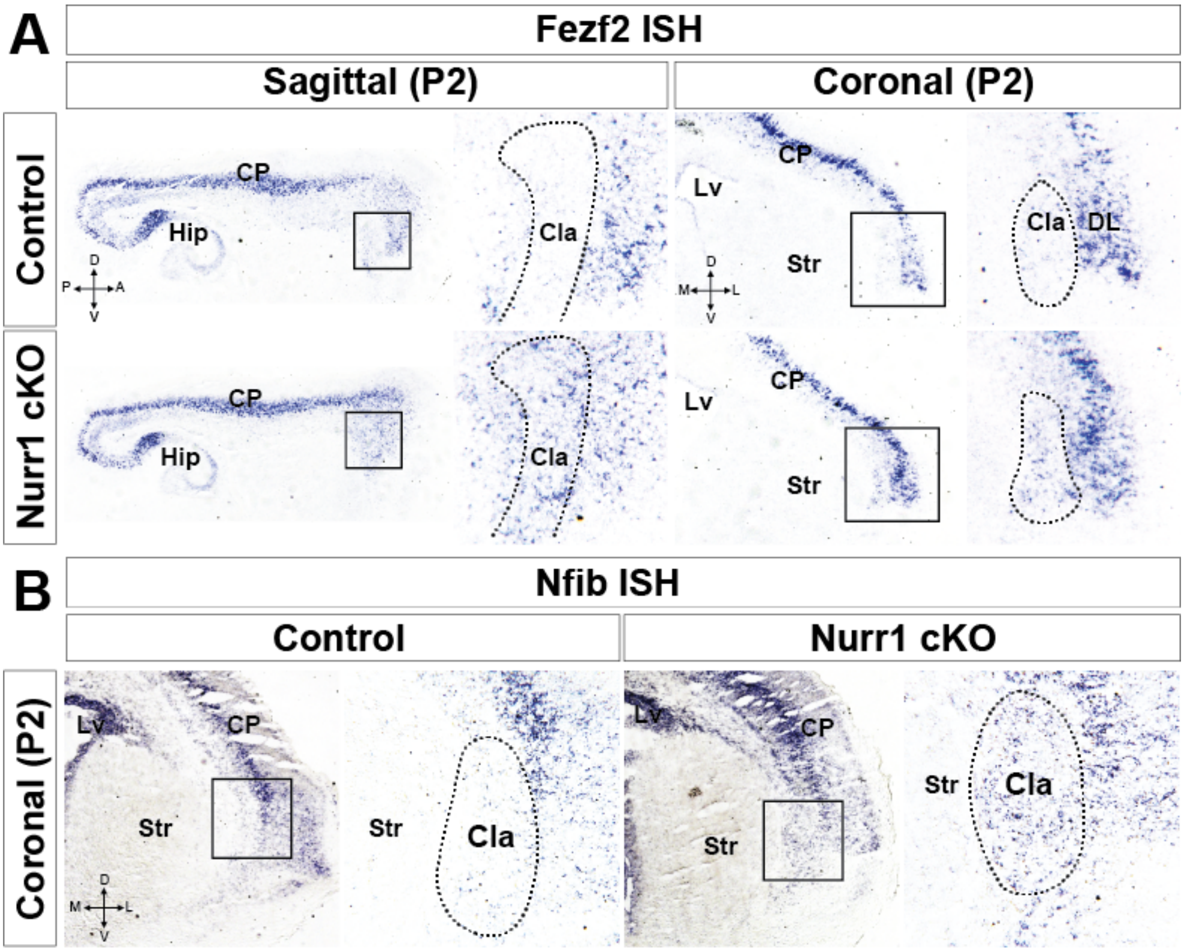


**Figure S4 Fezf2 and Nfib expression is ectopically detectable in claustrum of Nurr1 deficient brains.**

**(A, B)** ISH using the Fezf2 and Nfib probes on sagittal (**A**) and coronal (**B**) sections of control and Nurr1 deficient brains at P2. Fezf2 and Nfib expression is barely detected in CLA of normal brains but Fezf2+ and Nfib+ deeper layer (DL) neurons are populated in original CLA area of Nurr1 deficient brains.

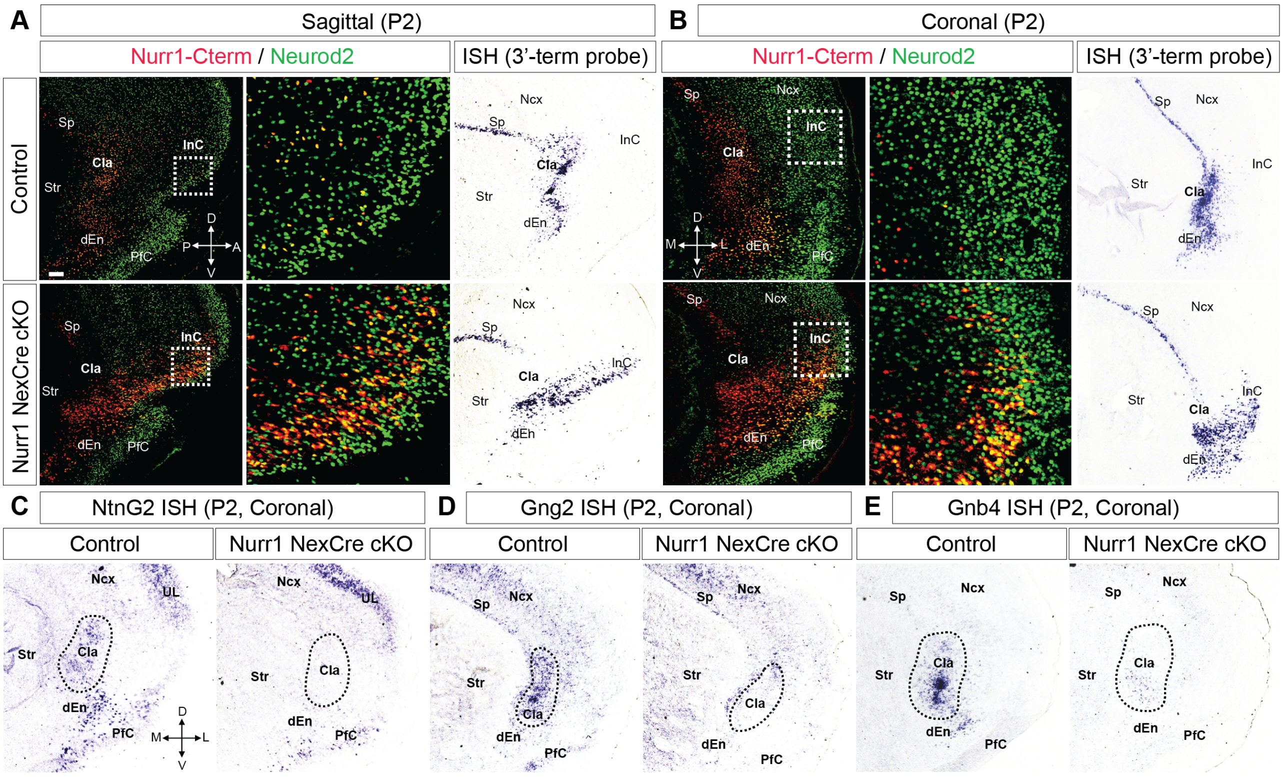


**Figure S5 The NexCre+ Nurr1 deficient claustral neurons recapitulate the same phenotypes as those of EmxCre+ mutant brains.**

**(A, B)** IF for Nurr1-Cterm (red) and Neurod2 (green) on sagittal (**A**) and coronal (**B**) sections of control and NexCre+ Nurr1 deficient brains at P2 shows that Nurr1+/Neurod2+ neurons aberrantly migrate into InC in NexCre+ mutant brains, mimicking the same phenotypes as in EmxCre+ Nurr1 deficient brains. ISH using the Nurr1 3'end probe on sagittal and coronal sections of control and NexCre+ mutant brains at P2 shows the same results consistent with IF staining. The magnification images show the enlarged views of boxed areas. Scale bar: 200 μm.

**(C-E)** ISH using the NtnG2 (**C**), Gng2 (**D**) and Gnb4 (**E**) probes on coronal sections of control and NexCre+ Nurr1 deficient brains at P2 shows that the expression of these genes is considerably downregulated in CLA and/or dEn cells of NexCre+ mutant brains, similar with EmxCre+ Nurr1 deficient brains. UL, upper layers of Ncx.

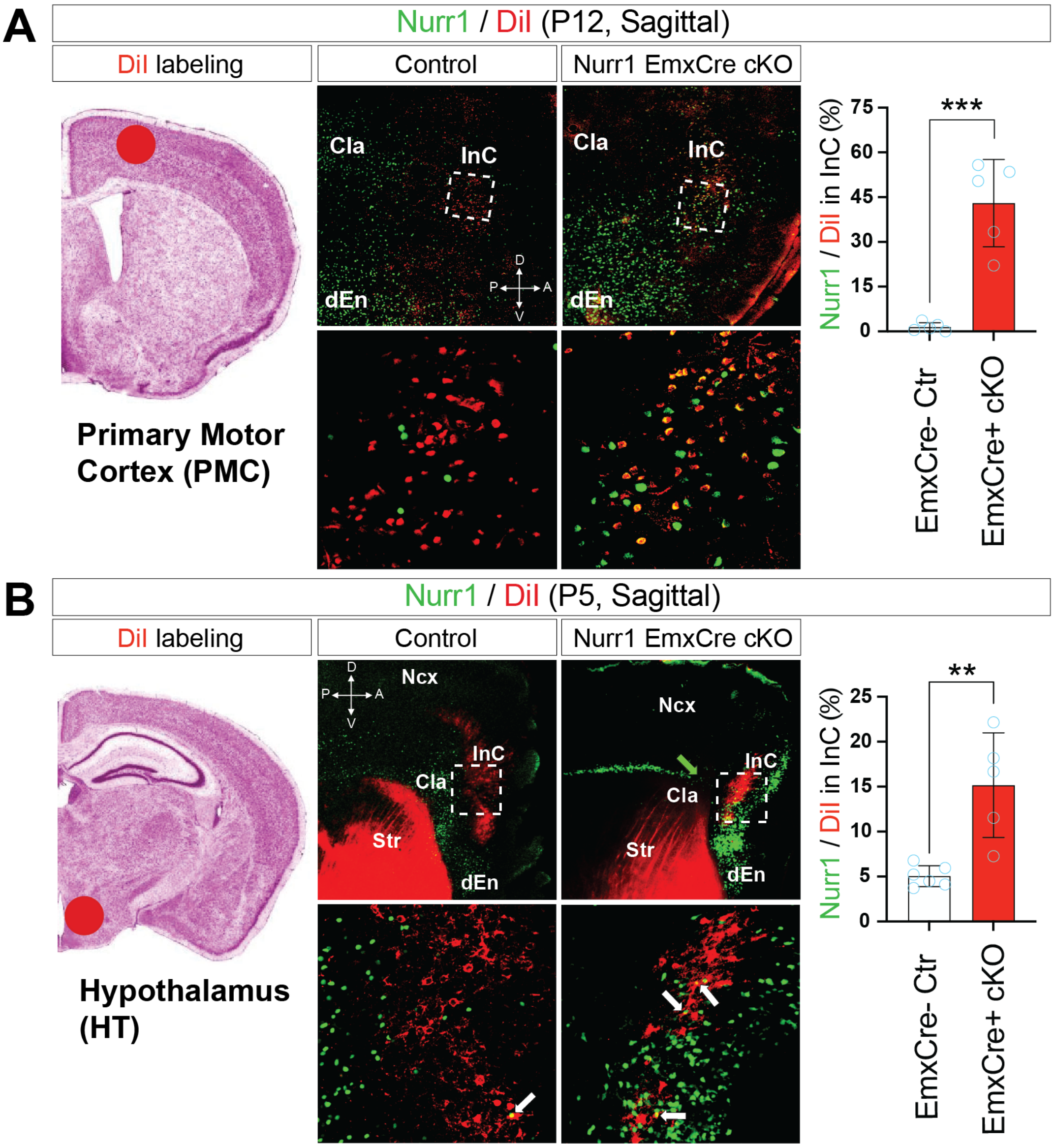


**Figure S6 Nurr1 deficient claustral neurons project ectopically along the axonal route of insular cortical neurons.**

**(A)** DiI crystals were placed into primary motor cortex (PMC) of P12 brains for axonal retrograde tracing. IF for Nurr1-Cterm (green) on brain sections carrying DiI shows that DiI+ signals traced from axonal terminals in PMC areas are mostly found in insular cortical cells, but barely colocalized with Nurr1+ CLA cells in control brains. In contrast, Nurr1 deficeint cells migrate into insular cortical area and ectopically colocalized with DiI+ signals. The ratio of Nurr1+/DiI+ cells relative to all DiI+ cells is ≈ 1.45% in control brains (n = 5), but significantly elevated up to ≈43.02% in Nurr1 deficeint brains ($p$ = 0.00023, ***; n = 5).

**(B)** DiI crystals were buried into hypothalamus (HT) of P5 brains for axonal retrograde tracing.

DiI+ somas traced from HT axonal terminals are also found in insular cortex neighboring to CLA in control brains, however, more Nurr1 deficeint cells were found colocalized with DiI+ signals in mutant brains. The green arrowhead indicates the CLA region of a Nurr1 deficient brain, and white arrowheads indicate Nurr1+/DiI+ cells. The ratio of Nurr1+/DiI+ cells is ≈5.05% in control brains (n = 6), but elevated to ≈15.16% in cKO brains ($p$ = 0.0023, **; n = 5).

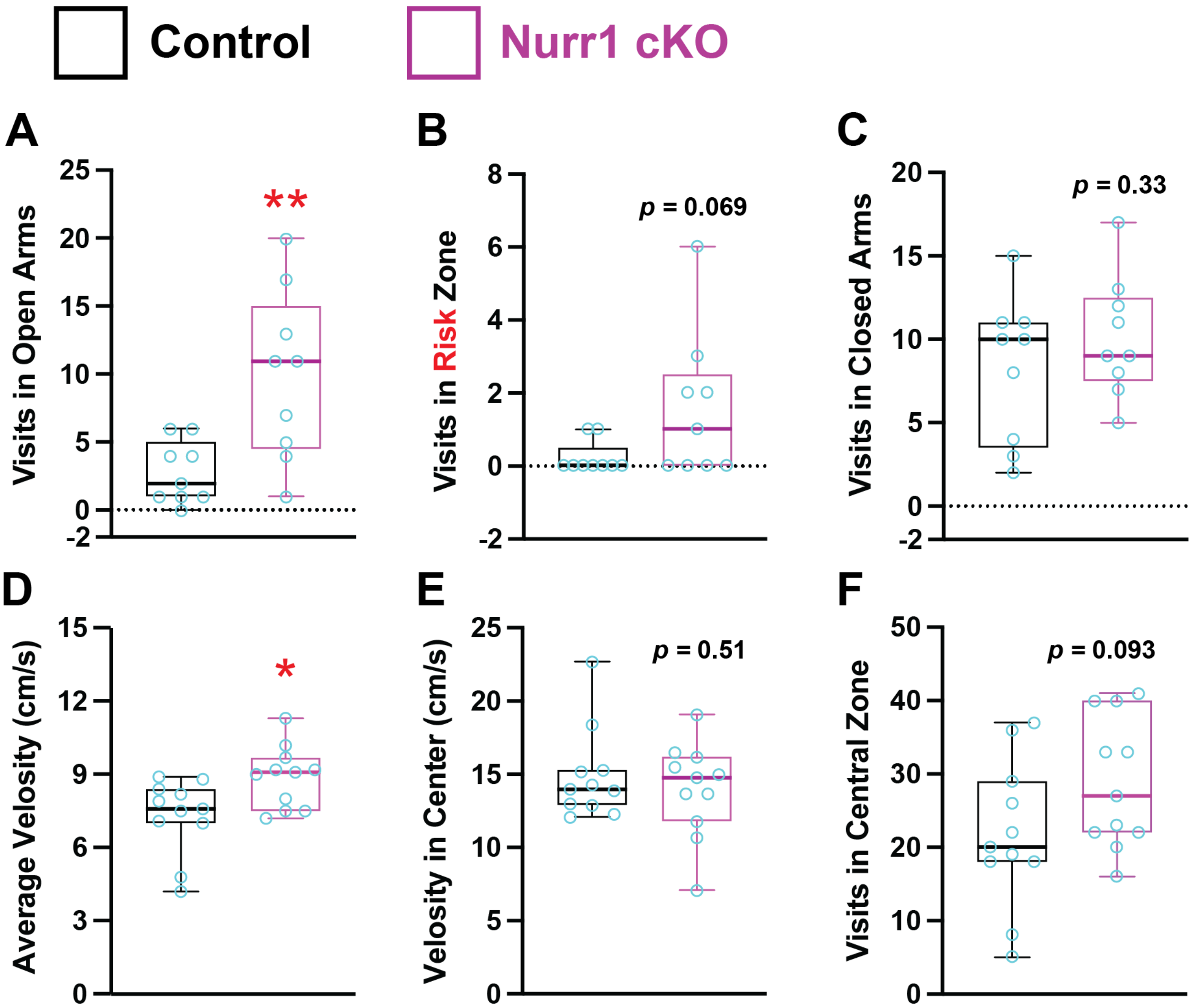


**Figure S7 Analysis of stress-elicited behaviors for Nurr1 deficient mice**

**(A-C)** The visit frequency in open arms, risk zones and closed arms in elevated plus maze. Nurr1 deficient mice visited the open arms (≈9.89, $p = 0.0054$, **; n = 9) and risk zones (≈1.56, $p = 0.069$, ns; n = 9) more often than control mice (open arms in controls: ≈2.78, n = 9) (risk zones in controls: ≈0.22, n = 9). Nurr1 deficient mice visited closed arms at a similar frequency (≈8.22, $p = 0.33$, ns; n = 9) with control mice (≈10.11, n = 9).

**(D)** The average velocity of mice in open field (OF) test: the velocity of Nurr1 deficient mice (≈8.90 cm/s, $p = 0.015$, *; n = 11) was mildly but significantly higher than control mice (≈7.31 cm/s, n = 11), suggesting the greater locomotion of Nurr1 deficient mice in OF.

**(E, F)** The velocity (**E**) and visit times (**F**) of mice in the central zone of OF. Nurr1 deficient mice exhibited similar velocity in the center (≈14.92 cm/s, $p = 0.51$, ns; n = 11) with control mice

(≈14.01 cm/s, n = 11). Notably, the animal velocity in the central zone is much higher than their average velocity irrespective of genotypes, indicating all animals were stressful in the central area of OF. Nurr1 deficient mice tended to visit the central area more frequently (≈28.82, n = 11) in comparison to control mice (≈21.64, n = 11), but not statistically significantly ($p$ = 0.093, ns).

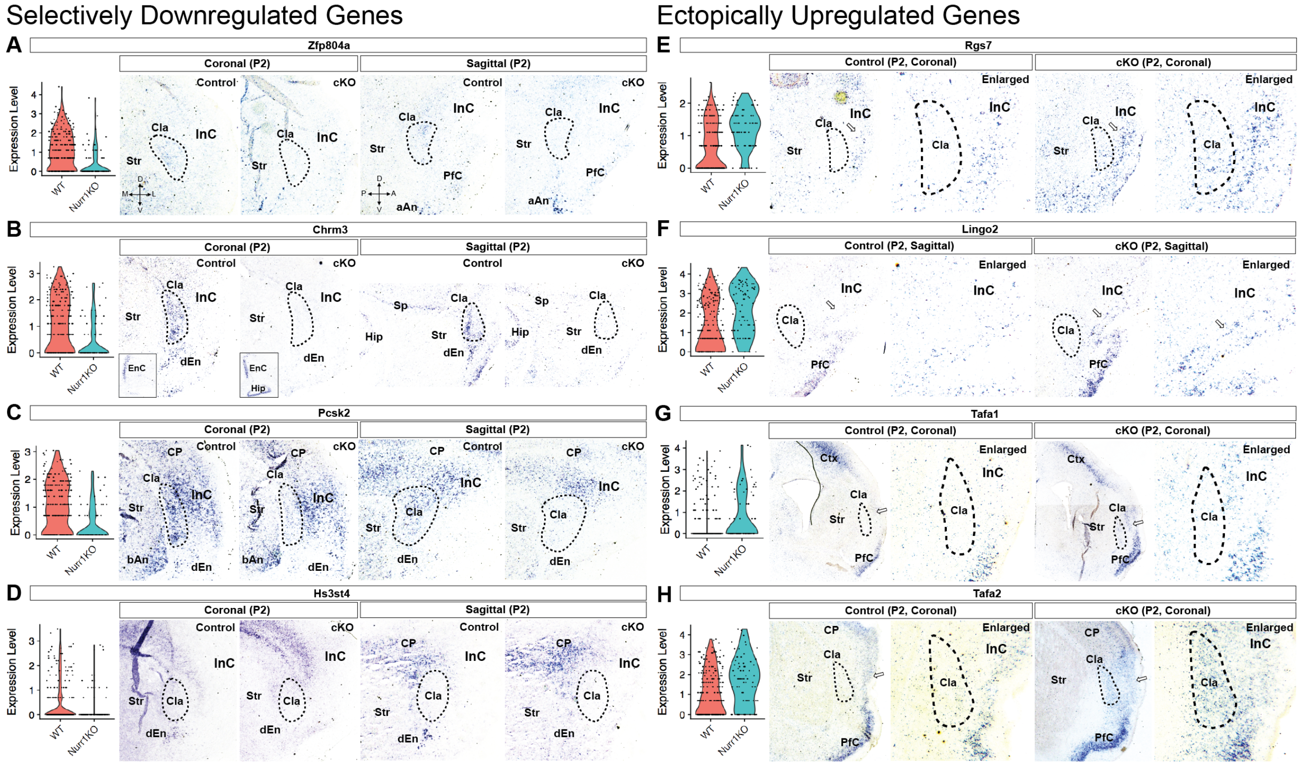


**Figure S8 ISH verification of misregulated genes in EmxCre+ Nurr1 deficient brains**

**(A-D)** ISH verification of downregulated genes in EmxCre+ Nurr1 deficient CLA and/or dEn cells. ISH using the Zfp804a (**A**), Chrm3 (**B**), Pcsk2 (**C**) and Hs3st4 (**D**) probes on coronal and sagittal sections of control and EmxCre+ mutant brains at P2. The violin plots of target genes (**A-H**) resulted from single cell mRNA sequencing are seated left to the ISH data. Zfp8104a expression is downregulated in Nurr1 deficient CLA cells but not in anterior amygdalar nuclei (aAn). Chrm3 expression is selectively downregulated in Nurr1 deficient CLA and dEn cells but not in entorhinal cortex (EnC) or Hip. Pcsk2 expression is downregulated in Nurr1 deficient CLA cells but not in basomedial amygdalar nuclei (bAn) and surrounding cortical neurons. Hs3st4 expression is selectively reduced in Nurr1 deficient dEn cells but not in surrounding DL neurons.

**(E-H)** ISH verification of ectopically upregulated genes in EmxCre+ Nurr1 deficient CLA cells. ISH using the Rgs7 (**E**), Lingo2 (**F**), Tafa1 (**G**) and Tafa2 (**H**) probes on coronal or sagittal sections of control and Nurr1 deficient brains at P2. The expression of Rgs7, Lingo2, Tafa1 and Tafa2 is barely detectable in CLA cells of control brains, however, their expression is upregulated in neighboring neocortex in Nurr1 deficient brains. The magnification views of CLA and InC are presented in the enlarged images.

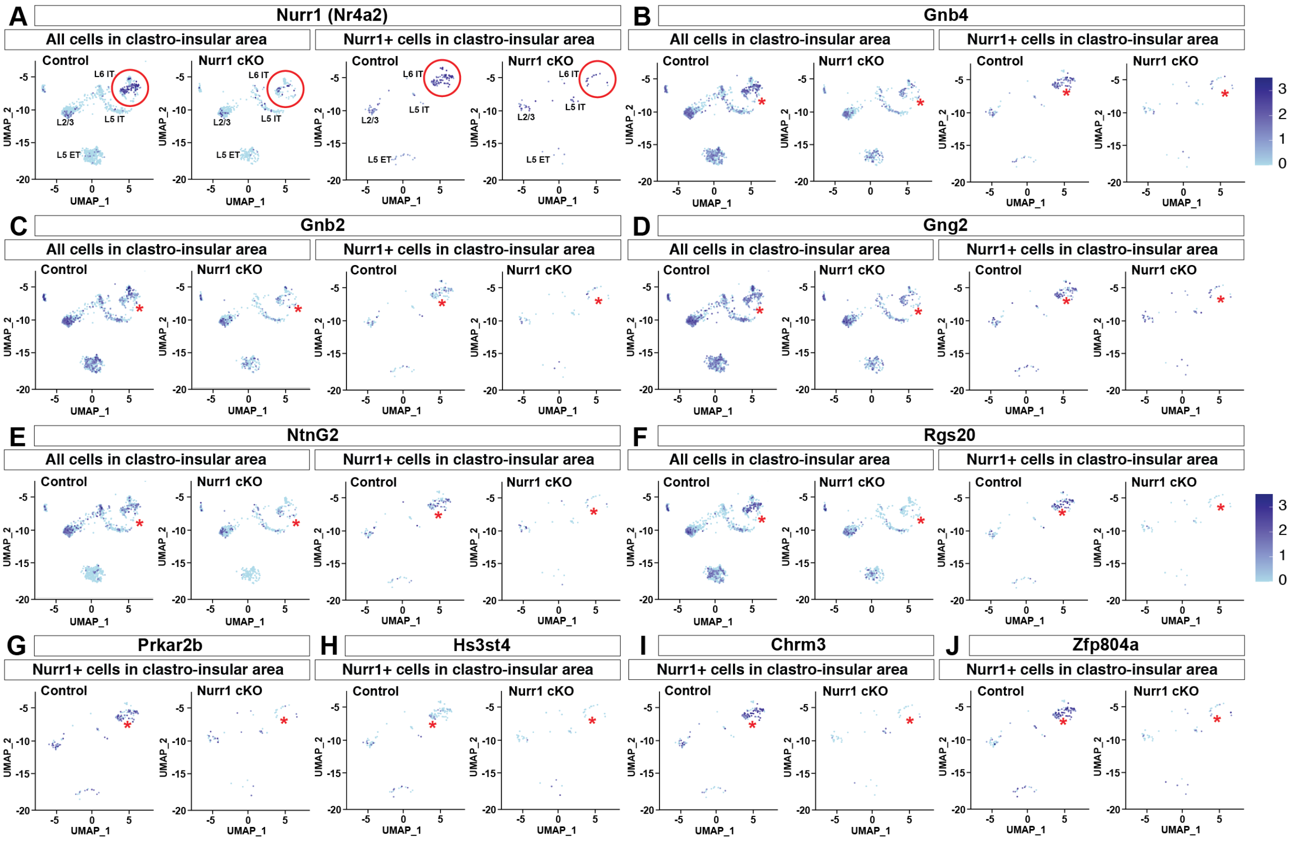


**Figure S9 UMAP dimensionality reduction of selected gene expression in control and Nurr1 deficient mice**

**(A-F)** UMAP analysis of single cell transcriptomes for Nurr1, Gnb4, Gnb2, Gng2, NtnG2 and Rgs20 in either all or Nurr1-expressing clastro-insular cells shows that these genes are downregulated in Nurr1+ cell populations of clastro-insular region of Nurr1 deficient brains. Nurr1 is predominately expressed in L6 cluster containing CLA and dEn cells (red circles in Nurr1 panel and asterisks in other panels). The expression level is marked from light blue (0, minimal) to dark blue (3, maximal) (the same in **G-J**). L2/3, layer 2 and 3 cells; L5 ET and IT, layer 5 extra- and intra-telencephalon projecting glutamatergic neurons; L6 IT, layer 6 intratelencephalon projecting glutamatergic neurons; L6b, layer 6b neurons (Sp).

**(G-J)** UMAP analysis for Prkar2b, Hs3st4, Chrm3 and Zfp804a in Nurr1-expressing cells shows that these genes are also downregulated in Nurr1+ cell populations of clastro-insular region in Nurr1 deficient mice.

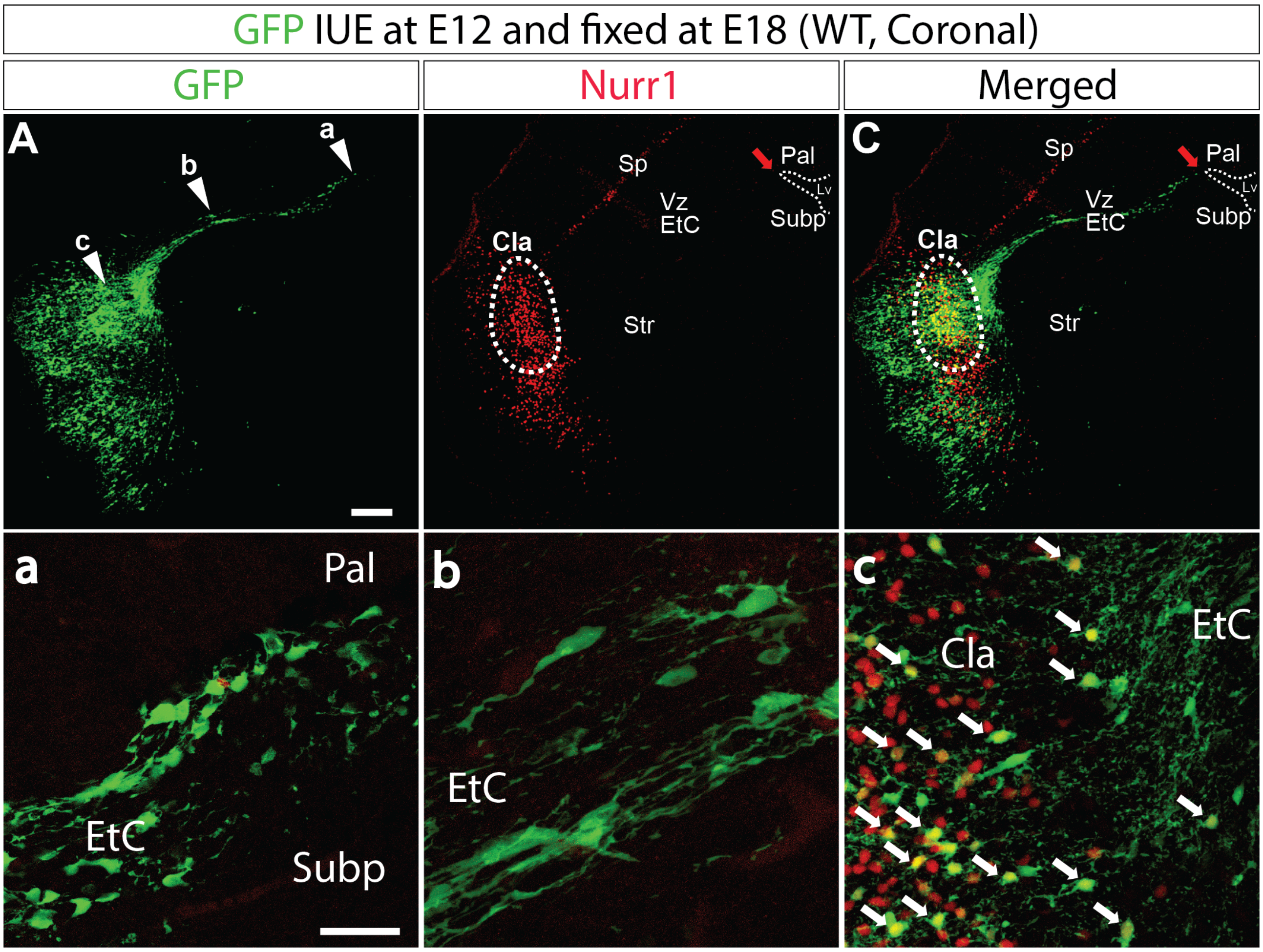


**Figure S10 GFP-electroporation at E12 in the progenitors at pallial-subpallial border targets the claustral cells.**

**(A-C)** *In vivo* electroporation (IUE) of GFP into wild type (WT) brains at E12 targeting neural progenitors at pallial-subpallial (Pal-subp) border illustrates a population of neurons migrate tangentially in the external capsule (EtC) towards lateral pallium. These cells later leave EtC, of which a major subpopulation become Nurr1+ CLA cells (yellow). Vz, ventricular zone. The dotted line circles indicate the CLA. The red arrowheads indicate the Pal-subp border. Scale bar: 200 µm.

**(a-c)** The enlarged views of indicated areas in (**A**). These immature cells originate from Pal-subp border (**a**) and migrated in EtC (**b**). CLA cells start to express Nurr1 intermediately after they leave EtC and reside at CLA area (**c**). Each white arrowhead indicates one or one cluster of GFP+/Nurr1+ cells. Scale bar: 50 µm.

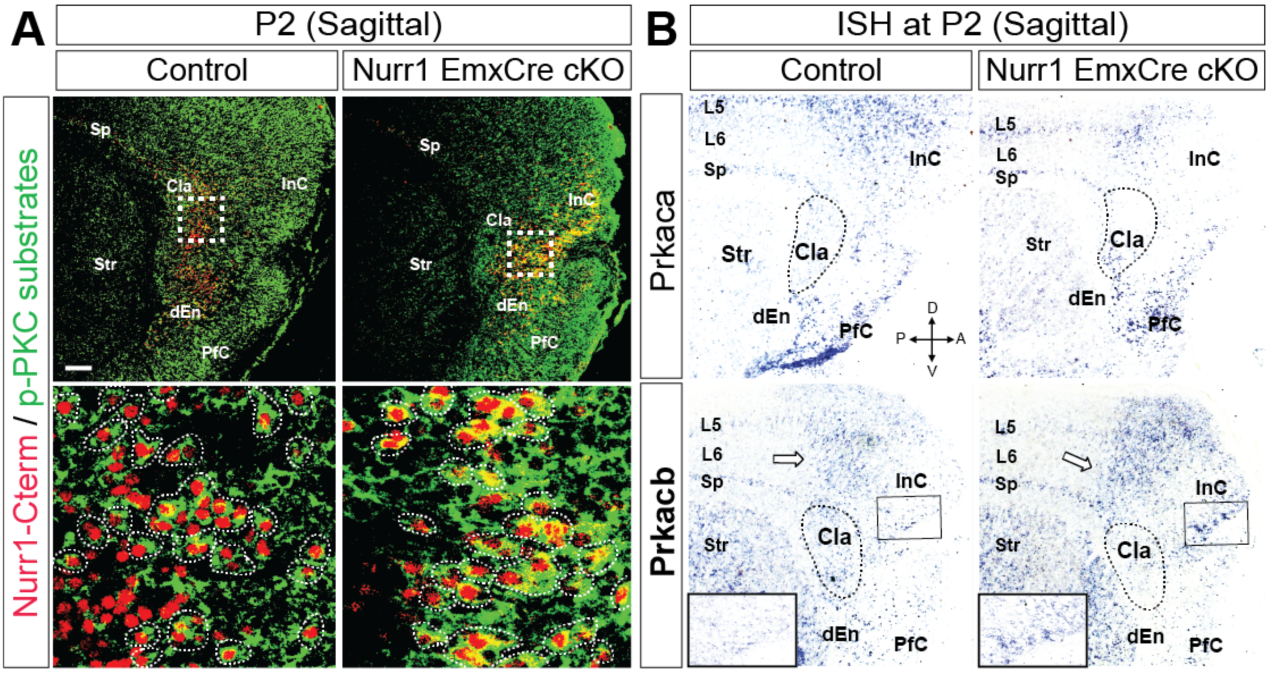


**Figure S11 One of PKA catalytic subunit, *Prkacb*, is ectopically upregulated in InC.**

**(A)** IF for Nurr1-Cterm (red) and phosphorylated substrates of PKC (green) on sagittal sections of control and Nurr1 deficient brains at P2 shows PKC is active in most Nurr1-expressing cells of both control and mutant brains. Each dotted line circle indicates one or one cluster of double positive cells. Scale bar: 200 µm.

**(B)** ISH using the Prkaca and Prkacb probes on sagittal sections of control and Nurr1 deficient brains at P2 shows that the expression of *Prkacb*, one of the PKA catalytic subunit, is ectopically activated in the InC of mutant brains. The hollow arrowheads indicate the borderlines of *Prkacb* positive signals, highly consistent with IF staining in Fig. 6E.

**Table S1 Oligos and Primers**

| Name: | Usage: | Sequence: 5' – 3' |
|---|---|---|
| Nurr1-Nterm-Fw | Nurr1 ISH for 5'end | TCCTCGCCTCAAGGAGCCAGCCCCG |
| Nurr1-Nterm-Rv | | AAGTGCGAACACCGTAGTGCTGACA |
| Nurr1-Cterm-Fw | Nurr1 ISH for 3'end | GTTAAAGAAGTGGTTCGCACGGACA |
| Nurr1-Cterm-Rv | | CTTAGAAAGGTAAGGTGTCCAGGAAA |
| Cux2-Fw | Cux2 ISH | GATGGAGACAGCCAGCCCCAGG |
| Cux2-Rv | | TTCAGAATTCCCACTCCAGGAC |
| Cdh13-Fw | Cdh13 ISH | TCGCTACTTATCAACTGTATGTGGA |
| Cdh13-Rv | | TGGGTCCTTGTAGATAGAGTACCTG |
| Fezf2-Fw | Fezf2 ISH | TCATGTGATGTCAGCTGAATGTAAA |
| Fezf2-Rv | | TGGAGTCCAGGTAGTTGAAGTAGTA |
| Nfib-Fw | Nfib ISH | TCAATGTATCAGAGCTTGTGAGAGT |
| Nfib-Rv | | AAGGGAATTAGTGACTGTAAGTGCT |
| NtnG2-Fw | NetrinG2 ISH | GAAGGATTATGTCAAGGTCAAAGTG |
| NtnG2-Rv | | CGATATTGGAGATGGCATAGAAGTA |
| Gnb4-Fw | Gnb4 ISH | ATATACAACCTAAAGACCCGAGAGG |
| Gnb4-Rv | | GAGAACAGAAAATATGGCACTCAAT |
| Gng2-Fw | Gng2 ISH | GGGGAAGCTGCTCTCTAACCAAGCC |
| Gng2-Rv | | GACAGCTTATCAGAGGGTATTTGAA |
| Rgs20-Fw | Rgs20 ISH | AGTAGGAACCGCTCTGACTAGTGTA |
| Rgs20-Rv | | CTGTTTTCTCAGCTAAGGACGTAAG |
| CTGF-Fw | CTGF (Ccn2) ISH | AGTTACCAATGACAATACCTTCTGC |
| CTGF-Rv | | CCACGGTAGTTAAAAACACAGATTT |
| S100a10-Fw | S100a10 ISH | ATGATGCTTACGTTTCACAGGTTTG |
| S100a10-Rv | | CCATTGGATTAAGTTTTCTCTCTCA |
| Zbtb20-Fw | Zbtb20 ISH | GACACATTCACTGACAAACTCTCAC |
| Zbtb20-Rv | | AGTCATAGTCATCTTCCATTTCCTG |
| Prkaca-Fw | Prkaca ISH | GAAGATCTTAGACAAGCAGAAGGTG |
| Prkaca-Rv | | ATAGTCGTCAAAGTTACTCGTGTCC |

| | | |
|---|---|---|
| Prkaca-Fw | Prkacb ISH | TGGATTGCTATTTATCAGAGAAAGG |
| Prkaca-Rv | | CTTACTATCTCACGGAGTGAAGAGC |
| Nurr1-Fw-XhoI | Generation of Nurr1 expression construct | TTTTTCTCGAGAGCCATGCCTT-GTGTTCAGGCGCAGTAT |
| Nurr1-Rv-NotI | | TTTTTGCGGCCGCTTA-GAAAGGTAAGGTGTCCAGGAA |
| Gnb4-Fw-XhoI | Generation of Gnb4 expression construct | TTTTTCTCGAGGATGAGCGAGCTGGAG-CAGCTGAGG |
| Gnb4-Rv-NotI | | TTTTTGCGGCCGCTCAATTCCAGAT-TCTAAGAAAACT |
| Gng2-Fw-XhoI | Generation of Gng2 expression construct | TTTTTCTCGAGACCATGGCCAG-CAACAACACCGCCA |
| Gng2-Rv-NotI | | TTTTTGCGGCCGCTTAAAGGATGGCG-CAGAAGAACT |
| Tafa1-Fw | Tafa1 ISH | CCTAAAATCCATTATCCCTGTCTTT |
| Tafa1-Rv | | TGCTGTAACTAAGAAATGAGTGCTG |
| Tafa2-Fw | Tafa2 ISH | TAACCCATTAACCACTCCTAAATCA |
| Tafa2-Rv | | ACAACATGTAGGCTTTCAGAGTTTC |
| Zfp804a-Fw | Zfp804a ISH | AAGAAACAATGAACACAACAGTGAA |
| Zfp804a-Rv | | GTTTGCTCTGAGTTCTTCTCGATAC |
| Chrm3-Fw | Chrm3 ISH | CCTTGTAGAAAAGGGGTTTATCAAT |
| Chrm3-Rv | | ATTTGTATGAAACTTGAACTGCACA |
| Hs3st4-Fw | Hs3st4 ISH | CTCTTCATGTGCACCCTGTC |
| Hs3st4-Rv | | GGGTCAAGAAAGGAGGGACA |
| Pcsk2-Fw | Pcsk2 ISH | GTGACTCGACCTTTATTTCTGTCAT |
| Pcsk2-Rv | | TCATAACAGGGAACTCTAAAACCAC |
| Lingo2-Fw | Lingo2 ISH | GTATACCTGACCCACCTTAACCTCT |
| Lingo2-Rv | | TTTTAAGGTCCAGGGAGAAAGTATT |
| Rgs7-Fw | Rgs7 ISH | AGATTGATCATCCTTGTATCTGAGC |
| Rgs7-Rv | | TCTCTTCTTGTCCACTTTAGCTTGT |
| Nurr1-GT-Fw | Nurr1 floxed genotyping | GCTGGAGCCAGAGTTGGAAG |

| | | |
|---|---|---|
| Nurr1-GT-Rv | | ATTCCTTGGAGACCTTCTCT |
| EmxCre-PrimerA | EmxCre genotyping | TCGATGCAACGAGTGATGAG |
| EmxCre-PrimerB | | TTCGGCTATACGTAACAGGG |
| EmxCre-PrimerC | | AAGGTGTGGTTCCAGAATCG |
| EmxCre-PrimerD | | CTCTCCACCAGAAGGCTGAG |
| NexCre-PrimerA | NexCre genotyping | CCGCATAACCAGTGAAACAG |
| NexCre-PrimerB | | GAGTCCTGGAATCAGTCTTTTTC |
| NexCre-PrimerC | | AGAATGTGGAGTAGGGTGAC |

**ISH,** *in situ* hybridization.

**Table S2 Primary antibodies**

| Primary Antibody | Species | Company | Cat # | Dilution | Usage | RRID |
|---|---|---|---|---|---|---|
| Nurr1 N terminal | Mouse | Santa Cruz | sc-376984 | 1:300 | IF | AB_2893391 |
| Nurr1 C terminal | Goat | R&D systems | AF2156 | 1:500 | IF | AB_2153894 |
| Nestin | Mouse | Merck | MAB353 | 1:800 | IF | AB_94911 |
| Zbtb20 | Rabbit | SIGMA | HPA016815 | 1:300 | IF | AB_1858947 |
| GFP | Rabbit | ThermoFisher | A-11122 | 1:500 | IF | AB_2307355 |
| GFP | Chicken | Abcam | Ab13970 | 1:1000 | IF | AB_300798 |
| Tle4 | Mouse | Santa Cruz | sc-365406 | 1:500 | IF | AB_10841582 |
| Darpp32 | Rabbit | Abcam | ab40801 | 1:500 | IF | AB_731843 |
| Cleaved Caspase 3 | Rabbit | Cell Signaling | 9579 | 1:500 | IF | AB_10897512 |
| Cytochrome P450 26b1 (Cyp26b1) | Mouse | Merck | MABS497 | 1: 500 | IF | unidentified |
| Neurod1 | Rabbit | Proteintech | 12081-1-AP | 1:1500 | IF | AB_2877823 |
| Rorβ | Mouse | R&D systems | PP-N7927-00 | 1:500 | IF | AB_1964364 |
| Phospho-(Ser/Thr) PKA substrates | Rabbit | Cell Signaling | 9621 | 1:800 | IF | AB_330304 |
| Phospho-PKC sub-strates | Rabbit | Cell Signaling | 6967 | 1:800 | IF | AB_10949977 |

**IF,** immunofluorescence.
